# Regulatory control circuits for stabilizing long-term anabolic product formation in yeast

**DOI:** 10.1101/2020.04.26.062273

**Authors:** Vasil D’Ambrosio, Eleonora Dore, Roberto Di Blasi, Marcel van den Broek, Suresh Sudarsan, Jolanda ter Horst, Francesca Ambri, Morten O.A. Sommer, Peter Rugbjerg, Jay. D Keasling, Robert Mans, Michael K. Jensen

**Affiliations:** Novo Nordisk Foundation Center for Biosustainability, Technical University of Denmark, Kgs. Lyngby, Denmark; Department of Biotechnology, Delft University of Technology, Delft, The Netherlands; Joint BioEnergy Institute, Emeryville, CA, USA; Biological Systems and Engineering Division, Lawrence Berkeley National Laboratory, Berkeley, CA, USA; Department of Chemical and Biomolecular Engineering, Department of Bioengineering, University of California, Berkeley, CA, USA; Center for Synthetic Biochemistry, Institute for Synthetic Biology, Shenzhen Institutes of Advanced Technologies, Shenzhen, China

## Abstract

Engineering living cells for production of chemicals, enzymes and therapeutics can burden cells due to use of limited native co-factor availability and/or expression burdens, totalling a fitness deficit compared to parental cells encoded through long evolutionary trajectories to maximise fitness. Ultimately, this discrepancy puts a selective pressure against fitness-burdened engineered cells under prolonged bioprocesses, and potentially leads to complete eradication of high-performing engineered cells at the population level. Here we present the mutation landscapes of fitness-burdened yeast cells engineered for vanillin-β-glucoside production. Next, we design synthetic control circuits based on transcriptome analysis and biosensors responsive to vanillin-β-glucoside pathway intermediates in order to stabilize vanillin-β-glucoside production over ∼55 generations in sequential passage experiments. Furthermore, using biosensors with two different modes of action we identify control circuits linking vanillin-β-glucoside pathway flux to various essential cellular functions, and demonstrate control circuits robustness and 92% higher vanillin-β-glucoside production, including 5-fold increase in total vanillin-β-glucoside pathway metabolite accumulation, in a fed-batch fermentation compared to vanillin-β-glucoside producing cells without control circuits.

## Introduction

To support sustainable and environmentally friendly production-processes, research focuses on the production of bio-based alternatives to petroleum-based production processes, made by engineered cell factories (Kruyer and Peralta-Yahya, 2017). However, in order to design commercially attractive bioprocesses for converting cheap renewable substrates into chemicals, proteins and fuels, innovative bioprocess technologies need to be developed and applied. For this purpose, online monitoring and sampling is essential in modern fermentation processes, in order to control and optimize bioreactor conditions for biobased production (Gomes et al., 2018). Usual parameters analysed online include pH, exhaust CO_2_, temperature, aeration, agitation, and dissolved oxygen. Likewise more advanced methods are continuously developed and applied for online fermentation monitoring for the immediate implementation of control actions, thereby dynamically improving overall bioprocess performance (Fazenda et al., 2013). However, most often, any of these parameters are mere proxies for evaluating the actual biocatalysis, i.e. the microbial production of a chemical or protein of interest, which is often analysed offline during or following completion of the actual fermentation (Gomes et al., 2018). The lack of techniques for facile online monitoring and control of formation of the product of interest limits the understanding and rational optimization of bioprocesses.

The lack of monitoring and control becomes critical as bioreactor volume and bioprocess duration increase, as biobased production over prolonged cultivation regimes is challenged by low-performing cells arising due to the inherent stochasticity of biological systems, and/or evolutionary drift away from energy-requiring heterologous anabolic reactions, and towards improved fitness (Rugbjerg and Sommer, 2019; Wang and Dunlop, 2019; Xiao et al., 2016). Nongenetic and genetic heterogeneity of microbial populations is largely acknowledged as evident from large differences in growth rate, resistance to stress, and regulatory circuit output of isogenic populations (Carlquist et al., 2012; Müller et al., 2010; Rugbjerg et al., 2018b). Even more so, in industrial high-cell density fed-batch bioprocesses, subpopulations exist that are many-fold different in these parameters from the population average (Wang and Dunlop, 2019; Xiao et al., 2016). These random variations arise because not all cells are exactly of the same size, some cells may have been mutated during prolonged seed trains and growth in large-scale cultivations, and nor do all cells have the same number of key components, incl., RNA polymerase, ribosomes, and other key factors governing the life of a cell (Elowitz et al., 2002; Müller et al., 2010; Rugbjerg et al., 2018b). This cell-to-cell physiological and genetic heterogeneity is acknowledged widespread, yet difficult to constrain for the purpose of biobased production from high-performing isoclonal populations called for in industrial bioprocesses. In fact, low-performing variants can account for up to 80-90% of the total cell population, yet produce less than half of the desired product (Xiao et al., 2016). Not only does this hamper the production yield, but it also results in inefficient nutrient utilization, and overall increased production costs. Thus, there is a strong motivation for developing new technologies that can monitor biocatalysis, and optimize bioprocesses by coupling detectable phenotypes to product formation in an efficient manner.

Small-molecule biosensors offer sensitive and real-time monitoring of product formation in microbial cell factories (David et al., 2016; Tao et al., 2017; Zhang et al., 2016), and are furthermore emerging as a promising technology for safeguarding high-performing productive cell factories from evolutionary drifting into low-performing ensembles during prolonged cultivations, as those often applied in industry (Rugbjerg et al., 2018b). Several groups have recently succeeded in engineering and applying small-molecule biosensors based on allosterically regulated bacterial transcription factors. These biosensors undergo conformational changes upon binding of specific intracellular ligands and can directly couple single-cell ligand accumulation to a change in reporter gene expression (e.g. fluorescence or antibiotic resistance). Biosensors detecting small-molecule accumulation can be employed for facile evaluation of subpopulation heterogeneity in diverse feedstock and bioreactor environments and supports prototyping optimal bioreactor conditions in relation to production of any candidate chemical for which a biosensor is available (Flachbart et al., 2019; Snoek et al., 2018; Xiao et al., 2016). Also, as the biosensors are genetically encoded, and couple product accumulation with gene expression output, these biosensors not only allow for monitoring of product accumulation at the single-cell level (diagnosis), but can also couple product accumulation to cellular growth and thereby enable selective growth advantage of high-performing subpopulation variants and/or shunting of competing metabolic pathway reactions (therapy). Indeed, in bacteria, the use of such biosensors has recently been applied to confer a growth advantage of high-performing non-genetic variants of fatty acid-producing cell factories compared with low-producers (Xiao et al., 2016), as well as for coupling fitness-burdening production of mevalonate with expression of essential genes by fine-tuned biosensor-driven control circuits and thereby stabilizing production for >90 cellular generations of cultivation (Rugbjerg et al., 2018b).

In this study, we evaluate the coupling of fitness-burdening product-formation to biosensor-controlled expression of ten different single essential genes covering four different metabolic functions of baker’s yeast *Saccharomyces cerevisiae*. We make use of *S. cerevisiae* engineered for production of industrially-relevant vanillin-β-glucoside, for which we first systematically analyse the metabolic conversions associated with fitness reduction, and thus determine evolutionary pressure points. This identifies the formation of a burdening vanillin-β-glucoside pathway intermediate as a main target for coupling production with expression of essential genes, which we validate by the use of biosensors with two different modes of action. Next, population-level genomics analysis verifies the evolutionary pressure points by which *S. cerevisiae* escapes fitness-burdening vanillin-β-glucoside production, and also illustrates robust genetically-encoded biosensor designs recalcitrant to evolutionary drifting over 50 generations. Ultimately, the best-performing biosensor-based control circuit is benchmarked with the parental vanillin-β-glucoside producing strain in a fed-batch fermentation in which strains with control circuits accumulate 5-fold higher total pathway intermediates, including 92% higher final vanillin-β-glucoside levels, compared to the parental strain.

## Results

### Selection and physiological characterization of production strain

To investigate the ability of genetically-encoded biosensors to stabilize production of ATP-requiring molecules in yeast, first a proof-of-concept production testbed was selected. Here we chose a *S. cerevisiae* cell factory engineered for the production of vanillin-β-glucoside (VG), previously reported to exhibit a 50% growth rate reduction compared to non-producing *S. cerevisiae* (Strucko et al., 2015), and for which we have recently engineered two relevant biosensors for VG pathway intermediates, namely PcaQ from *Sinorhizobium meliloti* for protocatechuic acid (PAC) detection, and VanR from *Caulobacter crescentus* for vanillic acid (VAC) detection (Ambri et al., 2020). The VG producing strain carries single-copy genomic integration of five heterologous genes required for synthesis of VG from the native shikimate pathway intermediate 3-dihydroshikimate (3-DHS), through the heterologous intermediates PAC, VAC, protocatechuic aldehyde (PAL), and vanillin (VAN)(Figure 1A-B).

**Figure 1.**
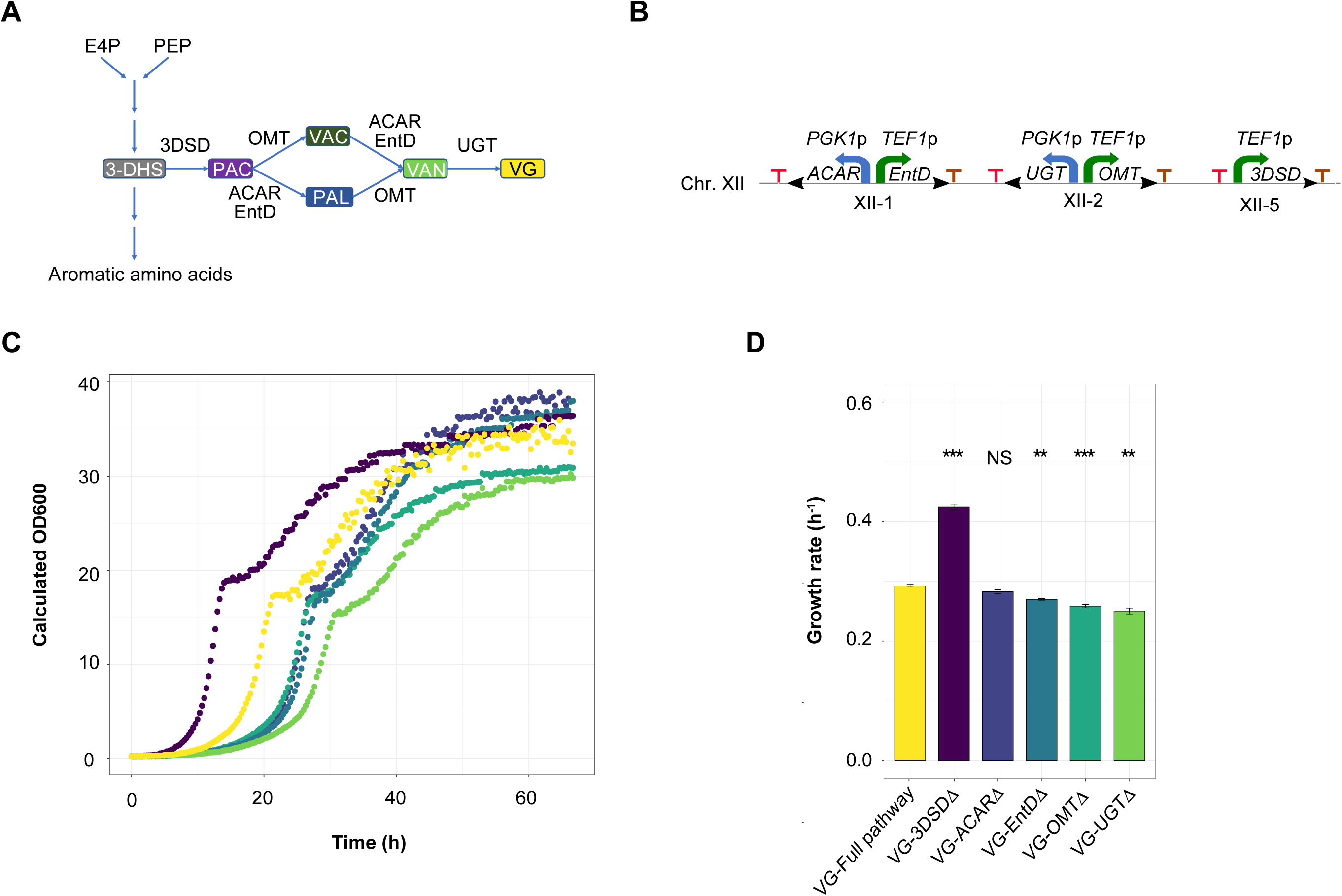
Identification of fitness-burdening reactions in the vanillin-β-glucoside biosynthetic pathway. (**A**) Schematic representation of the vanillin-β-glucoside biosynthetic pathway and the enzymes involved in vanillin-β-glucoside (VG) biosynthesis; 3DSD (3-dehydroshikimate dehydratase from *Podospora anserina*), hsOMT (O-methyltransferase from *Homo sapiens*), ACAR (aromatic carboxylic acid reductase from *Nocardia iowensis*), EntD (phosphopantetheine transferase from *Escherichia coli*), UGT (UDP-glycosyltransferase from *Arabidopsis thaliana*). (**B**) Schematic representation of biosynthetic pathway integration in EasyClone sites within chromosome XII. (**C**) Growth profile of single knockouts strains. Each curve is a representative of three (n=3) biological replicates. (**D**) Maximum specific growth rate observed over a 5-hours interval within the 67 hours cultivation. Single-knockout strains of genes of interest (GOI, indicated as VG-GOIΔ) are compared to the strain expressing the complete VG pathway (VG-Full pathway). Each bar represents the average of three (n=3) biological replicates with error bars representing mean ± standard deviation. Asterisks indicate statistical significance level as evaluated by t-test comparing growth rates for each of the different strains expressing a truncated pathway compared to the growth of the full VG pathway strain: NS = p > 0.5, ** = p <= 0.01, *** = p <= 0.001.

Next, in order to identify potential evolutionary pressure points for genetic instability of strains expressing the fitness-burdening VG pathway, we constructed single knockout variants of each of the five genes encoding the VG pathway enzymes, and tested the specific growth rates on synthetic medium permissive for VG formation of all resulting mutant strains compared to the parental VG producing strain with all five genes left intact (Figure 1C, Supplementary Figure S1). From this analysis we found that deletion of the *3DSD* gene (VG-3DSDΔ), encoding the enzyme catalysing the conversion from 3DHS to PAC, resulted in increased maximum specific growth rate and shortened lag phase compared to the full VG pathway strain, whereas deletion of any of the other genes in the pathway resulted in mutants with reduced growth rates compared to the parental VG pathway strain (Figure 1C-D). Furthermore, even though the growth rate noticeable differs only between the VG-3DSDΔ and VG pathway strain, we observed that when *ACAR, hsOMT* or *EntD* genes are deleted, the lag phase is significantly longer, suggesting that the introduction of the pathway redirects the flux of metabolites from the biosynthesis of aromatic amino acids to the pathway causing a delay in the exponential growth phase.

Combined, these results indicate that depletion of shikimate intermediates and/or build-up of VG pathway intermediate PAC impose a fitness burden to the cells, whereas the single loss-of-function of any other pathway gene results in growth disadvantage, corroborating earlier reports on vanillin toxicity and the response of *S. cerevisiae* to weak acids and lignin derivatives (Gu et al., 2019; Guo and Olsson, 2014; Hansen et al., 2009).

### Assessment of cell factory stability

Having evaluated the burden of the VG pathway on host fitness based on systematic pathway truncations, we next sought to investigate VG pathway metabolite profiles and pathway stability of the VG production strain. For this purpose, a serial passaging experiment consisting of seven sequential transfers (∼55 generations) was performed. Here cultures were initially grown in YPD medium and then transferred in synthetic medium (SM) (Verduyn et al., 1992) to acclimate them to the experimental conditions (Figure 2A). From this experiment we initially observed that following first transfer, in which cultures were diluted 1:100 and grown for 48 hours in fresh medium, the extracellular metabolite content was biased towards PAC, being the first intermediate of the VG pathway, accumulating to approx. 2 mM, while end-product VG accumulated to 0.78 mM (Figure 2B). This observation agrees with the original VG pathway study performed on this strain (Strucko et al., 2015). For each of the following transfers, cultures were further diluted 1:100 every 48 hours into fresh medium, thus initiating the next batch phase (Figure 2A). At the end of each transfer, samples were collected for quantification of all pathway intermediates and VG end-product formation in order to assess the total flux through the VG pathway in each batch phase of the passage regime.

**Figure 2.**
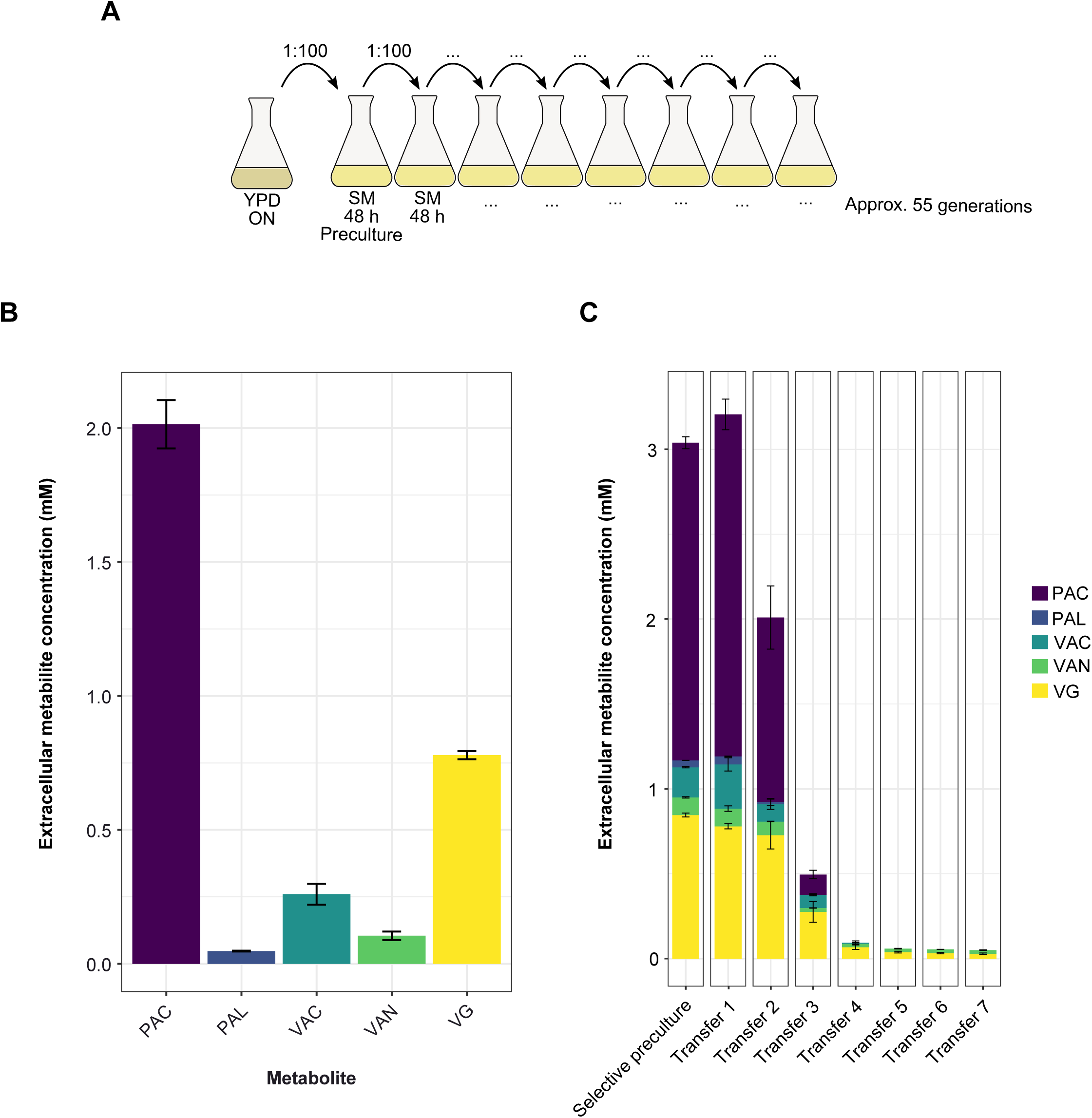
Assessing vanillin-β-glucoside biosynthetic pathway stability during prolonged cultivation using metabolite profiling. (**A**) Schematic layout of the sequential transfer regime of batch cultures. (**B**) HPLC analysis of the extracellular metabolite concentrations for the VG strain following the first sequential passage. (**C**) Extracellular metabolite concentrations for the VG strain after 48 h cultivations over seven sequential transfers. The bars represent the average of two (n=2) biological replicates. Error bars represent mean ± standard deviation from two (n=2) biological replicates.

Following the second transfer, a 46% drop in PAC productivity was observed compared to transfer 1, ultimately resulting in the inability to detect any extracellular PAC by the end of the fourth transfer. Similarly, VAC decreased by 60% and 97% by the end of the second and fourth transfer, respectively (Figure 2C). On the other hand, the decrease in VAN and VG concentrations was delayed compared to the other pathway intermediates. In the second transfer the extracellular concentration of VAN and VG only decreased by 24% and 6%, respectively. However, by the end of the fourth transfer VAN and VG concentrations decreased by 81.5 and 91.4%, respectively (Figure 2C), whereas by the end of the experiment VG had decreased by 98.5%.

Based on the accumulated evidence, 3DSD activity and/or depletion of shikimate intermediates appear to be causing the main fitness burden when expressing the VG pathway in yeast (Figures 1C-D and 2). For this reason, we sequenced the genomically integrated *3DSD* gene of 22 isolated single colonies from the fourth transfer of six parallel sequential batch cultures. From the sequencing analysis, we observed 7 premature stop codons, 5 SNPs, and 8 recombination events between the *3DSD* gene and either the gene encoding EntD or hsOMT, all controlled by the same *TEF1* promoter design and *CYC1* terminator (Figure 1B). The remaining 2 colonies sequenced did not have any mutations in the *3DSD* gene (Table 1).

**Table 1.** List of identified mutations in the *3DSD* gene. List of the identified mutations in the *3DSD* gene at the end of the fourth sequential passage from six parallel cultivations. Each entry represents the result from one of the 22 sequenced colonies.

| Mutations | Culture 1 | Culture 2 | Culture 3 | Culture 4 | Culture 5 | Culture 6 |
| --- | --- | --- | --- | --- | --- | --- |
| Stop codon | Q91X | W111X<br>S140X<br>W169X |  |  | Y92X<br>Y92X | Q124X |
| Homologous recombination | EntD<br>hsOMT | hsOMT | hsOMT | EntD<br>EntD | EntD | hsOMT |
| SNP |  | V243G | S11L<br>W80G<br>G35V | M280I |  |  |
| No mutations |  |  | Wild type |  |  | Wild type |

Taken together, these results suggest that the fitness-burdened parental VG producing strain is genetically unstable, and that the burden exerted by the pathway is predominantly alleviated via mutations in the first enzymatic step of the pathway encoded by *3DSD*, converting 3DHS into PAC. Moreover, the single knockout experiment shows that loss of any subsequent VG pathway gene in a strain with an intact *3DSD* gene negatively impacts the growth rate of the resulting truncated pathway designs (Figure 1B).

### Biosensor candidates and characterization

As already mentioned, control of population heterogeneity and stabilization of heterologous end-product formation has previously been established in bacteria by the use of small-molecule biosensors (Rugbjerg et al., 2018b; Xiao et al., 2016). In this study we wished to extend from this concept and assess the potential for control of production from heterologous pathways in eukaryotes making use of control circuits founded on prokaryotic small-molecule biosensors conditionally controlling expression of native genes essential to yeast. Moreover, we aimed to explore the hitherto unknown potential of engineering control circuits for stabilizing pathway intermediates instead of end product formation, thereby aiming to demonstrate the use of many more small-molecule biosensors for control circuit applications than possible if only considering biosensor-assisted control of end product formation.

For choice of biosensors we selected VanR from *Caulobacter crescentus* and PcaQ from *Sinorhizobium meliloti*, which we recently designed and applied as biosensors for VAC and PAC, respectively (Ambri et al., 2020; D’Ambrosio et al., 2020). Importantly, engineering control circuits by the use of either VanR or PcaQ, would enable testing of circuit performance founded on two different modes-of-action, potentially impacting the stability of the control circuits. Mechanistically, VanR is a transcriptional repressor which, in the absence of VAC, prevents transcription by binding to VanR operator sites (VanO) in promoters and thus confers sterical hindrance of RNA polymerase activity (Gitzinger et al., 2012). In the presence of VAC, VanR undergoes a conformational change, decreasing its affinity to VanO, upon which transcription can start (Jain, 2015). For the PAC biosensor, the transcriptional activator PcaQ, a LysR-type transcriptional regulator, constitutively binds PcaO operator sites in gene promoters and induces transcription of the output gene in the presence of PAC (Ambri et al., 2020; Fernandez-López et al., 2015). Regarding the design of the genetically-encoded biosensors, the VAC biosensor is composed of a bi-directional system where VanR is expressed under the control of the constitutive *PGK1* promoter while the output gene is controlled by a synthetic VanO-containing *TEF1* promoter (Ambri et al., 2020), supporting an operational range spanning >2 orders of magnitude of VAC concentrations, including the range of metabolite concentration observed in the parental VG strain (Figure 2, and Supplementary Figure S2). For the PAC biosensor, PcaQ is expressed from the strong constitutive *TDH3* promoter, while the output gene is controlled by a truncated PcaO-containing *CYC1* promoter (209 bp) with low activity in the absence of PAC, and a high dynamic output range (Ambri et al., 2020).

To characterize VanR and PcaQ biosensors for sensing VAC and PAC concentrations, respectively, we first introduced VanR or PcaQ together with either GFP-expressing VanO- or PcaO-containing reporter promoters in both the VG production strain as well as in the non-producing VG-3DSDΔ strain, and measured fluorescence outputs. Here, the introduction of VanR and PcaQ in the VG strain resulted in increases in GFP read-outs of 2.7- and 3.5-fold, respectively, compared to their expression in the VG-3DSDΔ strain (Figure 3), confirming that the biosensors are able to discriminate between VG-producing and non-producing strains.

**Figure 3.**
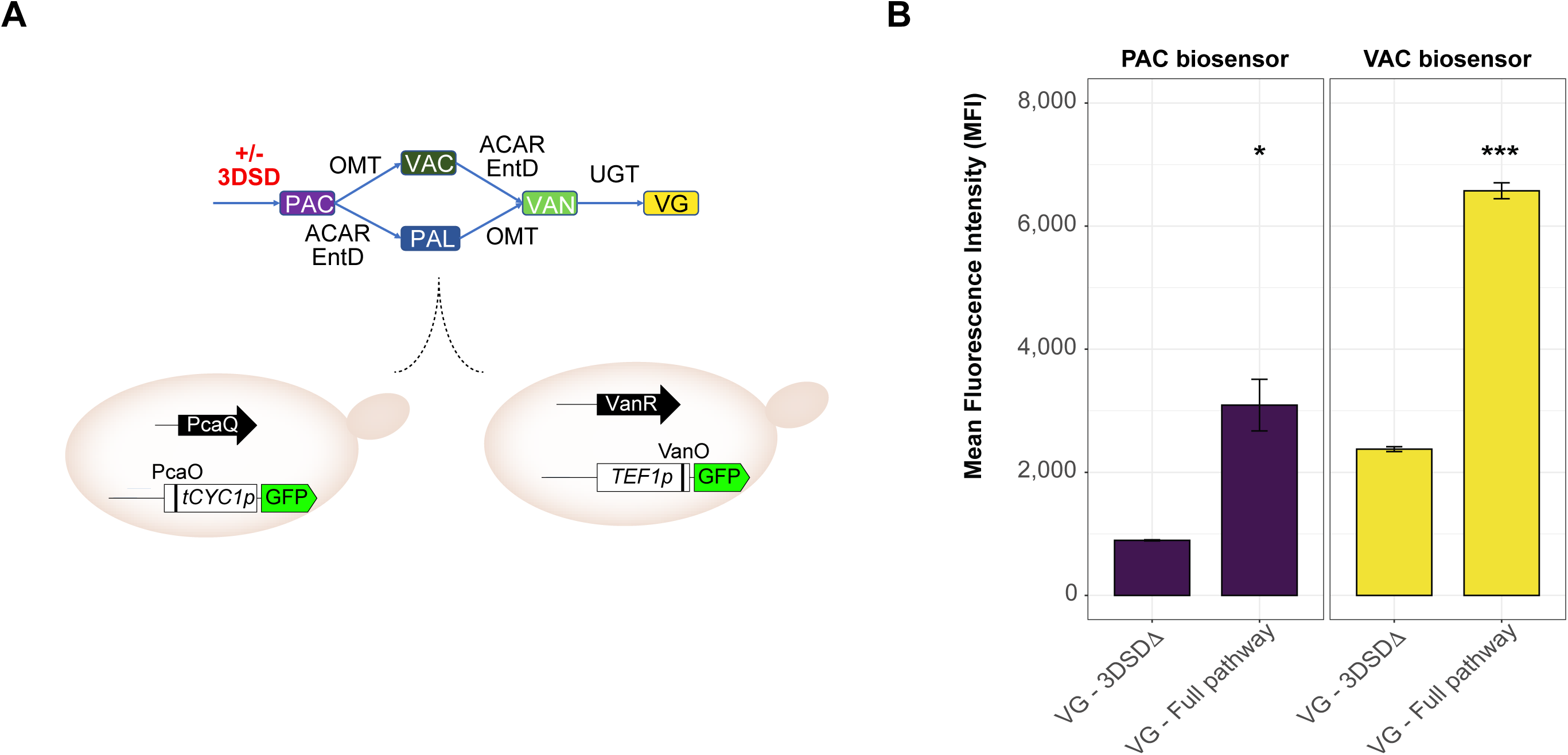
Biosensor design and *in vivo* validation. (**A**) VAC and PAC biosensor designs are tested in the complete (VG - Full pathway strain) or broken (VG-3DSDΔ) pathway strain. (**B**) Mean fluorescence intensity (MFI) of VanR and PcaQ biosensor designs in the vanillin-β-glucoside producing strains (VG-Full pathway) and non-producing (VG-3DSDΔ) strains. The values represent the average MFI of three (n=3) biological replicates. Error bars represent mean ± standard deviation from three (n=3) biological replicates. Asterisks indicate statistical significance level as evaluated by *t*-test comparing average MFIs for the VG-3DSDΔ strains expressing a truncated pathway compared to the MFIs of the full VG pathway strain: * = p <= 0.05, *** = p <= 0.001.

### Evaluation of control circuit designs

Spontaneous mutants with disruptive mutations in the *3DSD* gene, and therefore without the ability to produce VG and its biosynthetic intermediates, have a growth advantage compared to the strains harbouring the full VG pathway (Figure 1C-D). Eventually, such differences in growth rate and lag phase will allow for complete population take-over of non-producer cells during prolonged cultivations. To extend the productive life-span of a parental VG strain, we next sought to make use of the validated VAC and PAC biosensors for engineering control circuits in which mutants losing productivity will be subject to reduced fitness or even complete growth retardation. To do so, we decided to couple the expression of essential genes to the presence of VG pathway metabolites PAC and VAC, similar to previously reported studies in *E. coli* (Rugbjerg et al., 2018b; Xiao et al., 2016). Ideally, for such control circuits, only strains carrying a functional product pathway would be able to conditionally induce transcription of essential genes to a level sufficient for growth.

To select essential genes for control circuit designs, we performed a systematic analysis of candidate essential genes in yeast. From the total list of 5,188 validated open reading frames of the *S. cerevisiae* genome (Figure 4A)(Cherry et al., 2012), we initially selected four main classes of biosynthetic reactions (aiming to limit growth) that are not part of central carbon metabolism, namely 1) nucleotide metabolism, 2) cofactor/vitamin metabolism, 3) lipid metabolism, and 4) amino acid metabolism, containing a total of 325 unique genes (Figure 4A, Supplementary Figure S3, Supplementary Table S4). This list was further refined to focus on 3-4 selected metabolic pathways from each of the four classes (e.g. cysteine biosynthesis from homocysteine) totalling 110 unique genes, of which we omitted metabolic reactions catalysed by multiple gene products (e.g. *ADE5*/*ADE7*), bringing the gene list to 68 candidates (Figure 4A). Of special attention, it should be noted that for selection of genes involved in amino acid biosynthesis, we initially made a complete list of the abundance of amino acids in the heterologous proteins of the VG pathway and compared this to the average amino acid composition of yeast biomass (Lange and Heijnen, 2001). For example, glutamine accounts for only 3.98% of the amino acids in the VG pathway, whereas leucine is the most abundant accounting for 10.71% (Supplementary Table S5). For comparison, the same analysis for the average composition of yeast biomass, revealed that glutamine accounts for 7.75% of yeast biomass, and leucine for 8.03% (Supplementary Table S5), from which we hypothesized that a possible limitation in glutamine biosynthesis could effectively limit growth while having minimal impact on VG productivity. Finally, we decided to include *ARO2* as a candidate gene since it is involved downstream of the shikimate pathway (Gottardi et al., 2017), and for which limited expression could therefore help to accumulate more shikimate intermediate and thus boost VG pathway flux. To further refine the selection criteria to an operational number for control circuit testing, we focused our attention on 56 genes described as essential or causing auxotrophy (Cherry et al., 2012), and then only selected genes with an average expression level higher than the VanO-containing *TEF1* promoter in the absence of VAC (OFF) across a wide range of yeast growth rates (i.e. 0.02-0.33 h^-1^) in glucose-limited aerobic chemostats (Regenberg et al., 2006). Because of the low OFF state and dynamic range previously reported for the PcaQ biosensor design (Ambri et al., 2020), essential gene candidates with average expression within the VanR dynamic output range were also considered relevant for PAC control circuits founded on PcaQ. Finally, from this list of 34 genes, we manually selected 10 genes, where either accumulation or depletion of its respective substrate and product due to altered expression levels would not cause toxic effects (e.g. increased mutagenesis), as our candidate list of essential genes for control circuit testing (Figure 4A-B).

**Figure 4.**
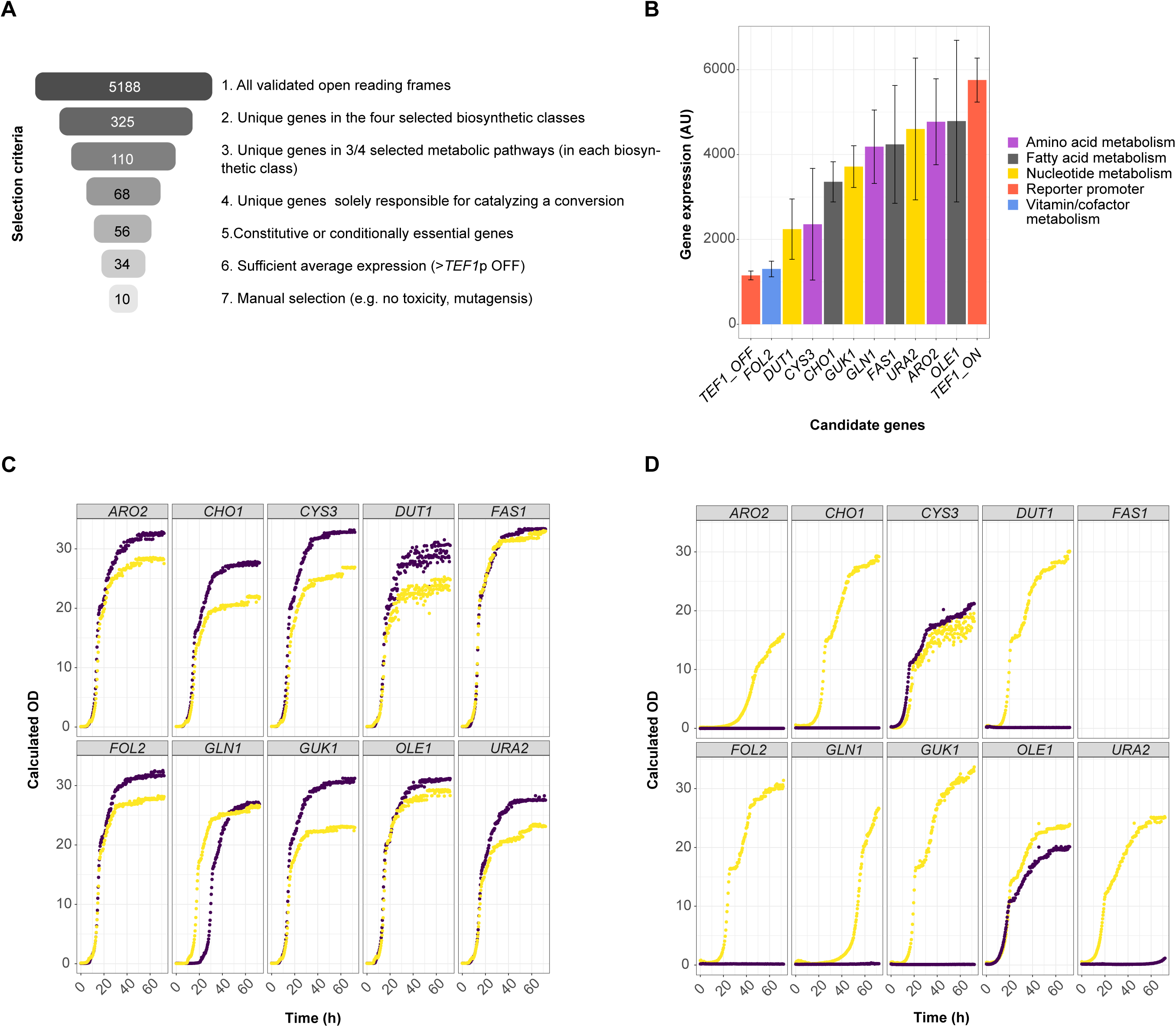
Selection of essential genes and testing of VAC and PAC control circuits. (**A**) Schematic illustration of the various filtering criteria applied for essential gene selection. The number of genes passing each of the seven filtering steps are indicated in the grey-coloured bars, and the steps filtering criteria indicated to the right. (**B**) Native expression levels of essential genes selected for promoter replacement. Genes were selected for different metabolic functions: nucleotide metabolism (yellow), fatty acid metabolism (black), amino acid metabolism (purple) and vitamin/cofactor metabolism (blue). (**C**) Growth profile of wild type CEN.PK strains carrying the VAC control circuit design in the presence (yellow) or absence (purple) of 2mM vanillic acid. (**D**) Growth profile of VG pathway strains (yellow) and VG-3DSDΔ strains (purple) carrying the PAC control circuit design. Each curve is representative of a minimum of two biological replicates. It was not possible to identify a colony having the *FAS1* promoter replaced by the PAC control circuit.

To test the performance of the control circuits, we initially introduced the VanR biosensor design in a prototrophic CEN.PK113-7D background strain by individually replacing the genomic locus, containing the first 150bp upstream of the first ATG of each of the 10 selected essential genes, with the synthetic construct (Supplementary Figure S4A). A similar approach was adopted for PAC control circuits, with the exception that PcaQ and the synthetic PcaO-containing *CYC1* promoter were introduced into both the parental VG producing strain and the non-productive VG-3DSDΔ strains (Supplementary Figure S4B). Following successful CRISPR-mediated promoter replacements, the strains were tested for conditional or improved growth upon external feeding of VAC or internal formation of PAC in VG-producing strains compared to VG-3DSDΔ strains for the VAC and PAC control circuits, respectively.

For strains expressing the VAC control circuits, cells were grown in SM for 24 hours, then diluted into fresh medium in the presence or absence of VAC, and growth was subsequently monitored for 72 hours. From this, it was evident that when coupled to the glutamine synthase encoded by *GLN1*, the VAC control circuit enabled shortening of the initial lag phase by approx. 10 hours in the presence of VAC, while the specific growth rate was not significantly altered between the different conditions (Figure 4C, Supplementary Figure S5). For the other 9 tested genes no VAC-dependent growth effects were observed.

Next, we assessed the growth when PcaQ was used to control the expression of the same set of 10 essential genes expressed from the PcaO-containing truncated *CYC1* promoter (Figure 4D, Supplementary Figure S6). Here, with the exception of *OLE1* and *CYS3*, and to a lesser extent also *URA2*, all control circuits had completely abolished growth of non-producing VG-3DSDΔ strains compared to VG-producing strains over 72 hours of cultivation (Figure 4D).

Taken together, this demonstrates that metabolic pathways involved in four different metabolic functions can be used for designing control circuits, and suggests that the low OFF state supported by the PcaO-containing truncated *CYC1* enables a broader range of essential genes covering a larger basal expression amplitude to be used, compared to control circuits founded on the repressor-type transcriptional regulator VanR controlling the synthetic VanO-containing *TEF1* promoter.

### Production stability of strains expressing control circuits

By design, control circuits founded on genetically-encoded biosensors controlling the expression of essential genes may themselves be pressure points for fitness-burdened cells to escape heterologous anabolic product formation. In order to investigate the ability of the engineered control circuits for stably maintaining product formation over long cultivation regimes, we next introduced the VAC and PAC control circuit in the VG production strain, and repeated the batch cultivation experiment with seven sequential passages, spanning a total of approx. 55 generations (Figure 2A). Again, after each transfer the extracellular concentration of metabolites was assessed by HPLC.

As determined by HPLC analysis, the parental VG strain maintained its productive lifespan for VG and all VG-pathway intermediates for one passage following the selective preculture (approx. 14 generations), yet with a >91% loss in VG productivity following four batch transfers, and a 98.4% loss of VG productivity by the end of the sequential passage experiment (Figure 2C, Figure 5A). In comparison, for the strains armed with VAC control circuits one culture showed a gradual decrease in VG productivity, yet maintaining approx. 70% VG productivity at the end of the experiment (0.73 mM to 0.52 mM VG), whereas the second culture had completely lost VG production by the end of the experiment (Figure 5B), indicating that the fitness-burdened VG producing strain can still escape the VAC control circuit and/or mutate *3DSD*.

**Figure 5.**
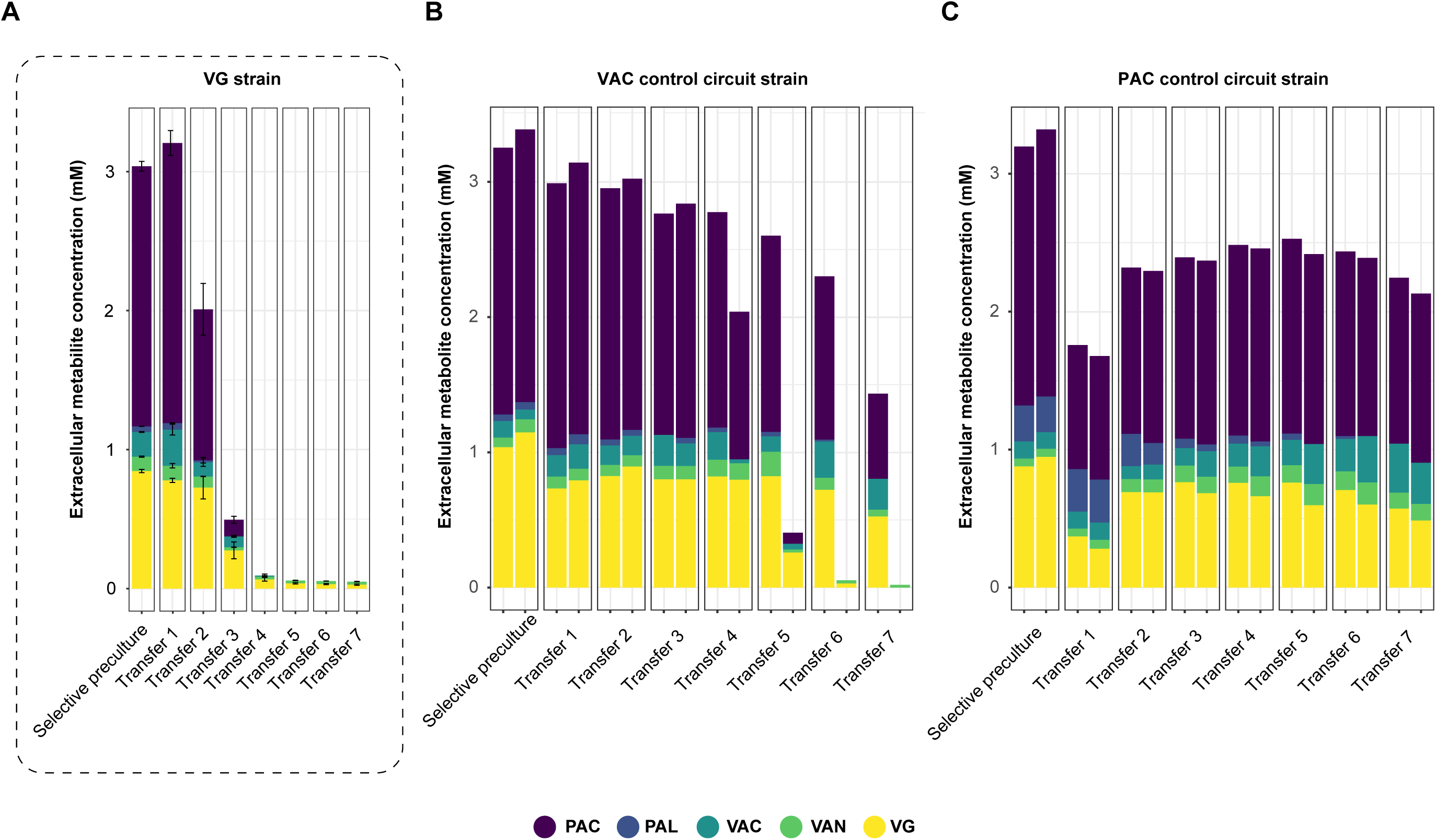
Assessing vanillin-β-glucoside biosynthetic pathway stability in biosensor control circuit strains. (**A**) Extracellular metabolite concentrations for the VG strain after 48 h cultivations over seven sequential transfers (Insert from Figure 2C). (**B**-**C**) HPLC analysis of the extracellular metabolites for strains harboring control circuits based on VanR (A) or PcaQ (B) during serial passage experiments. For (B-C) each bar represents total accumulated extracellular metabolites of the VG biosynthetic pathway based on one (n=1) biological replicate. For (B-C) each bar represents one biological replicate.

For strains expressing the PAC control circuit, we repeatedly observed >50% decrease in PAC and VG productivity from the selective preculture to transfer 1 (Figure 5C). Yet following the first transfer, productivity increased immediately during the following batch cultivations, ending the passage experiment with an increase in VG productivity of 54.5% and 72.2% for the two biological replicates, respectively (Figure 5C). Furthermore, all the VG intermediates, with the exception of PAL, were produced at higher concentrations by the end of the passage experiment. More precisely, both biological replicates presented an approx. 30% increase in PAC levels and approximately twice as much VAN compared to the first transfer. However, the biggest increase was observed for VAC where the two biological replicates increased productivity by 188% and 137%, respectively (Figure 5C).

Taken together, this time-resolved experiment demonstrates that control circuits coupling essential *GLN1* expression with accumulation of VAC and PAC, founded on the VanR transcriptional repressor and the transcriptional activator PcaQ, respectively, can enable extension of the productive lifespan of VG pathway, and in the case of PAC control circuit both pathway intermediates and final VG product formation is increased.

### Population-level sequencing to assess integrity of control circuits

Based on the results obtained from the batch cultivations of cells with and without control circuits (Figure 2, Figure 5), we next sought to determine whether i) the VanR-based control circuit mutated together with the VG pathway, and ii) if the cells expressing the PcaQ-based control circuits included mutations required to restore the growth rate and product formation following the first transfer (Figure 5). For this purpose, we whole-genome sequenced populations following the fourth and the seventh sequential culture for strains carrying the VanR-based control circuit and the third and the seventh culture for the strains with the PcaQ-based control of *GLN1* expression and we compared the results with their corresponding parental strains.

We mapped the whole-genome sequencing reads against the CEN.PK113-7D genome as a reference to assess genome coverage, base-calling and mutations. First, for the cultivation of the strains expressing the VAC control circuit, the replicate maintaining approx. 70% VG productivity following transfer 4 had no mutations identified, yet at transfer 7, a mutation localized in the VanO-containing *TEF1* promoter, driving the expression of the *3DSD* gene, was identified (Figure 6A). This mutation was observed in 23% of the reads from this population, and localized 18 bases downstream of the transcription start site of the *TEF1* promoter (Figure 6A). Moreover, in this replicate culture, two additional nonsynonymous mutations, encoding Q91P and P208S, were identified in 13% and 15% of the reads, respectively (Supplementary figure S7A-B). For the second replicate culture expressing the VAC control circuit, with a complete loss of VG productivity by the end of the cultivation, a single non-synonymous mutation in *3DSD*, encoding P208R, was observed in 63% of the reads by the end of the fourth culture (Figure 6B). Furthermore, approx. 40% of the reads showed a targeted 30 bp deletion in the *TEF1* promoter exactly covering the VanO sequence (Figure 6C). For the same replicate, at transfer 7 where no VG production was observed (Figure 5B), the P208R mutation in the coding region of the *3DSD* gene was present in 98% of the reads, and reads carrying the VanO site deletion increased to approx. 65% of the total population (Figure 6B-C). Finally, from the whole-genome sequence analysis of the PcaQ-stabilized strains no mutations were observed in the *3DSD* gene or the PcaQ biosensor design (Supplementary Figure S8).

**Figure 6.**
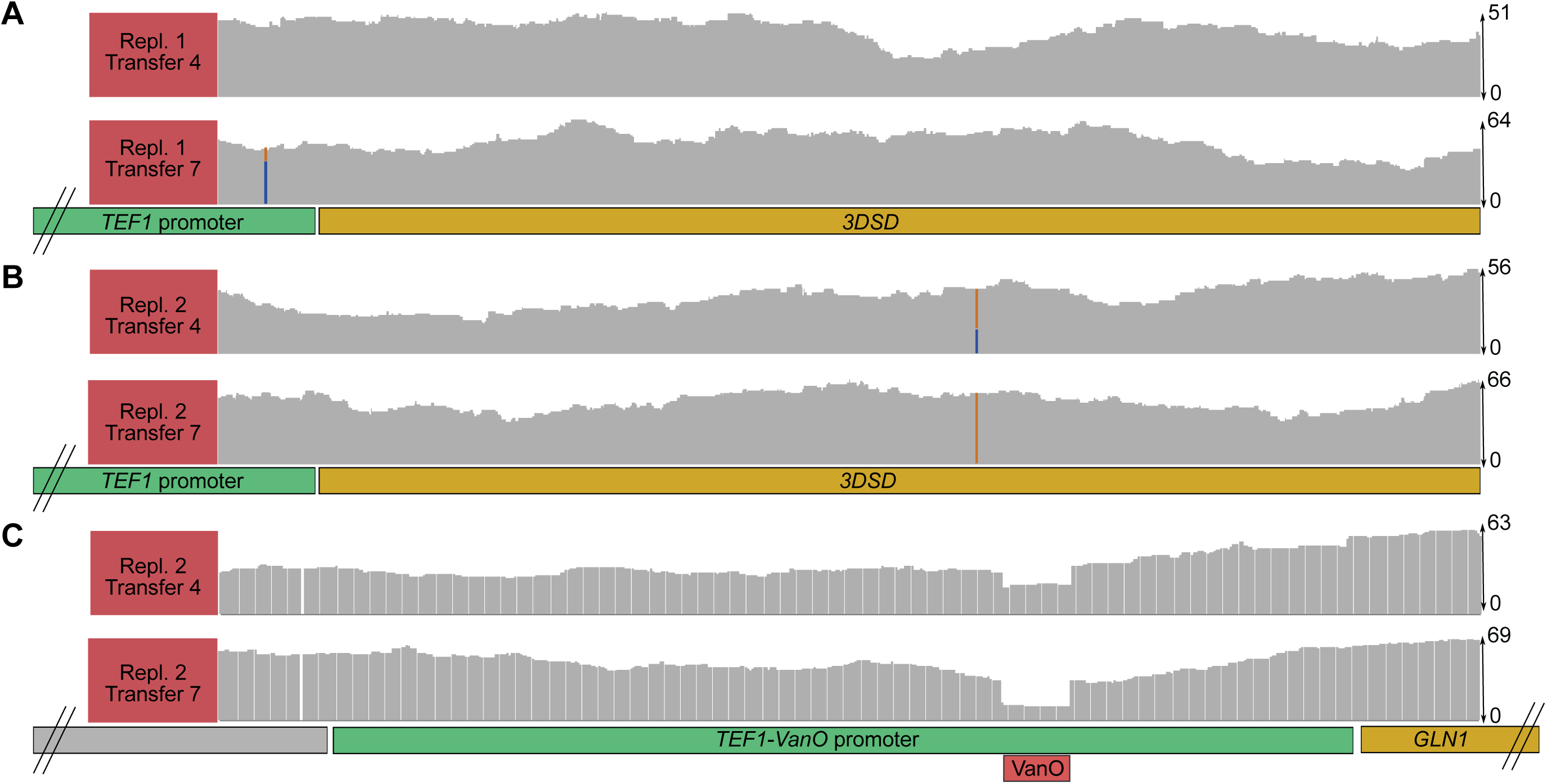
Whole-genome population level sequencing identifies the error-modes of VAC control circuits. (**A-B**) Genome-viewer zooms on the *3DSD* gene of the two replicate VAC control circuit strains at transfer 4 and 7 (see also Figure 5A). The blue and orange bars represent the ratio of wild-type (blue) and mutated (orange) sequence reads aligned to *3DSD*. Below the bar plots for read coverage, the genomic layout of the *3DSD* gene expression unit is indicated with the *TEF1* presented in green and the *3DSD* gene in yellow. (**C**) Genome-viewer zoom on the synthetic VanO-containing *TEF1* promoter of the VAC control circuit strain (replicate 2, Figure 5A). The VanO operator site is marked with a red box. Below the bar plot for read coverage, the genomic layout of the *GLN1* gene expression unit is indicated with the *TEF1-VanO* promoter presented in green and the *GLN1* gene in yellow. For (A-C) read coverage assembled to the CEN.PK113-7D reference genome is indicated to the right.

Taken together, these results show that, at population level, both the VAC control circuit and *3DSD* are mutated for yeast to escape fitness-burdened VG production. Furthermore, for the strains expressing the VAC control circuit, these results suggest that the *3DSD* gene is likely to mutate before the synthetic *TEF1* promoter controlling the expression of the *GLN1* gene. On the contrary, strains expressing the PAC control circuit, founded on the transcriptional activator PcaQ and the minimalized synthetic *CYC1* promoter, enables robust expansion of the productive lifespan of the VG pathway, without any observed genetic perturbations.

### Industrial scale-up mimic

Based on the successful application of the control circuits for coupling VG pathway stability to essential glutamine biosynthesis via the *GLN1* gene during sequential shake flask experiments, we tested the VAC and PAC control circuit designs in a downscaled fed-batch process in order to test the control circuit under industry-relevant conditions. First, to mimic the preceding seed train, we grew pre-cultures of strains with and without the VAC or PAC control circuit under non-selective conditions in rich medium. Following 24 hours, the pre-cultures were used to inoculate the first seed-train culture in a culture tube with a 1:100 inoculation ratio. Following another 48 hours, 40 µL of this first seed-train culture was transferred to a 4-mL shake flask culture, and after another 48 hours this culture was used to inoculate a 400-mL bioreactor. This seed train from 40 µL to 400 mL represents the same number of generations encountered in industrial seed trains starting from 1L and ending at 10,000 L (Fu et al., 2014).

The bioreactor culture started with a batch phase using 20 g/L of glucose and after carbon depletion, indicated by a rapid rise in dissolved oxygen levels and drop of off-gas CO_2_, the fed-batch phase was started (Figure 7, and Supplementary Figure S9A-C). During the fed-batch, a concentrated solution of feed medium was used, and the growth rate was set to 0.05h^-1^. Samples were taken at defined time-points (0, 20, 25, 44, 50, 68, 74 and 92 hours) during the batch and fed-batch phases, and metabolite consumption and formation was determined off-line via HPLC analysis (Figure 7, Supplementary Figure S10). The cultures were stopped when the working volume in the bioreactor reached 900 mL. For the parental VG strain, the batch phase stopped after 40 hours, during which approx. 1.4 mmol of VG pathway metabolites were formed from the initial 20 g/L of glucose (Figure 7). Similarly, the batch phase of the strain expressing VAC control circuit terminated after 40 hours, yet with 1.9 mmol of VG pathway metabolites accumulated (Figure 7). In contrast, the strain expressing the PAC control circuit grew slower, and the batch phase only finished after 55 hours, during which approx. 1.3 mmol VG pathway metabolites accumulated (Figure 7). These results are in agreement with the observed drop in productivity of the parental VG strain after the second transfer in the sequential batch culture experiment (Figure 2B), suggesting that a fraction of parental VG producing cells in the inoculum of the bioreactor already exhibited loss of productivity. Following the initial batch phase, the impact of the control circuits became even more apparent in the fed-batch phase. Here, the strain expressing the PAC control circuit outperformed both the parental VG strain and the strain expressing the VAC control circuit by producing 11 mmol of VG pathway metabolites compared to approx. 2.2 mmol and 8.9 mmol of VG pathway metabolites for the VG and VAC control circuit strains, respectively (Figure 7). Importantly, even though the strains expressing control circuits accumulate 4-5-fold higher amounts of total VG pathway metabolites compared to the parental VG strain, these strains also accumulate higher amounts of the VG end-product. Indeed, for the strains expressing the VAC and PAC control circuits, at the end of the fermentation, a total of 2.34 mmol and 2.61 mmol of VG, respectively, is produced, compared to 1.36 mmol of VG produced by the parental VG strain, representing 72% and 92% improvement in production (Figure 7). However, for the strains expressing the control circuits, there is also a substantial increase in the accumulation of VG pathway intermediates. For instance, and as also expected for control circuits founded on a PAC biosensor, in the strain expressing the PAC control circuit 6.8 mmol of PAC is accumulated, which is approx. 15 times the amount produced by the parental VG strain, and ∼40% more compared to the VAC control circuit strain. Likewise, the strain expressing the VAC control circuit accumulates more VAC (1.5 mmol) than both the parental VG strain and the PAC control circuit strain (0.3 mmol and 0.5 mmol, respectively) (Figure 7). Furthermore, in the parental VG strains PAC accumulates until the measurement at 50 hours, following which the PAC levels drastically decrease, corroborating the findings from the batch cultivations (Figure 2). Similarly, PAL, which is almost entirely converted to VAN in the parental VG strains, accumulates to 0.14 and 1.05 mmol at the end of the cultivation in the strains expressing the VAC and PAC control circuit, while for VAN the parental VG strain accumulate 0.02 mmol compared to 0.05 and 0.09 mmol for the strains expressing the VAC and PAC control circuits, respectively (Figure 7).

**Figure 7.**
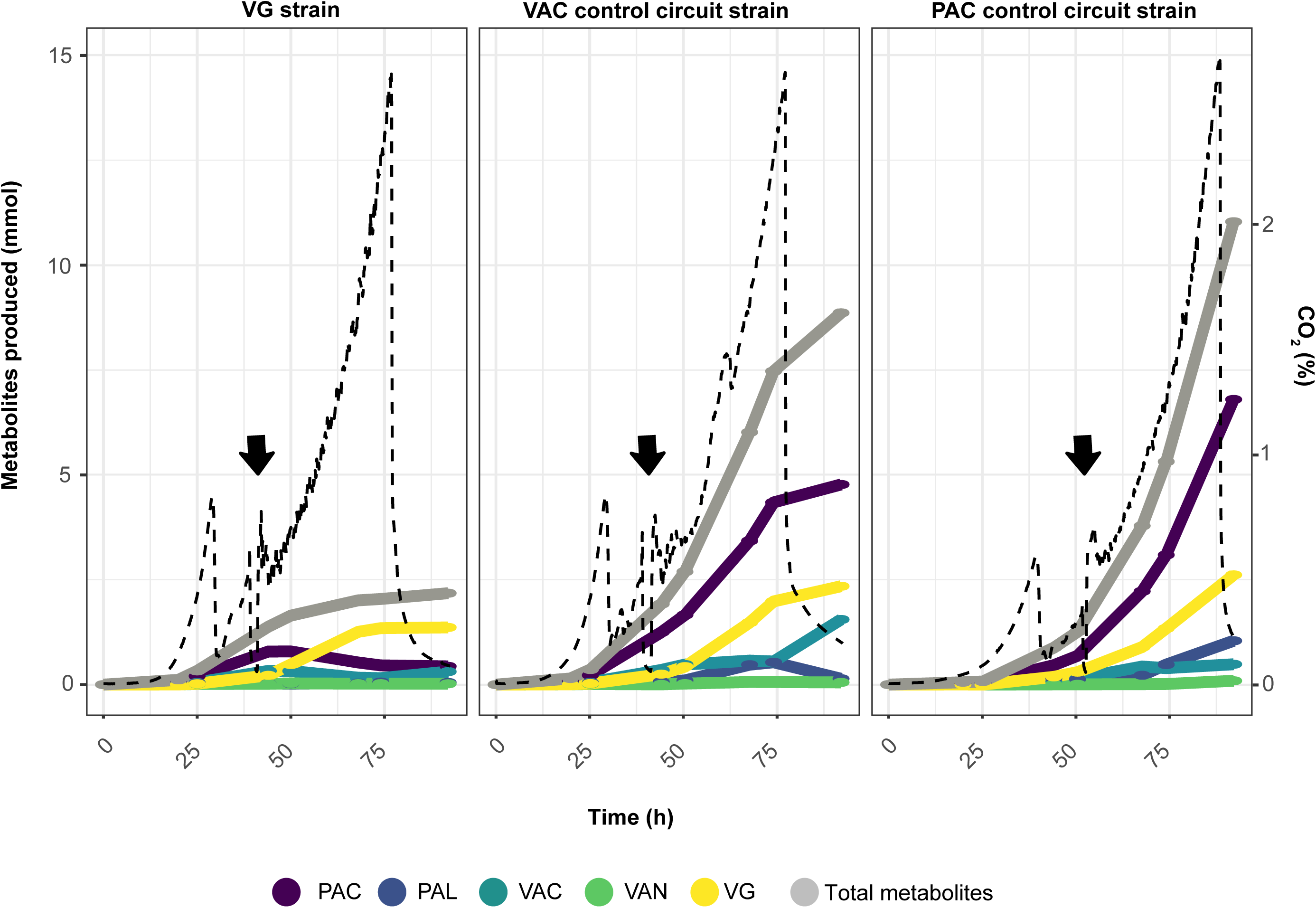
Fed-batch bioprocess for parental VG strain and strains expressing VAC and PAC control circuits. Metabolite and growth characterization of the parental VG strain (left panel), and strains expressing VAC (middle panel) and PAC (right panel) control circuits during the fed-batch bioprocess. Metabolite production is expressed in mmol of VG pathway metabolites produced during the cultivation. The values were corrected for the volume of medium removed for HPLC sampling and represent one (n=1) biological replicate. Data for a second biological replicate can be found in Supplementary Figure S10. Dashed black lines represent off-gas CO_2_ (percent), while black arrows represent the beginning of the fed-batch phase. Coloured lines indicate accumulation of individual VG pathway metabolites during the cultivation (Time, h) according to colour coding indicated at the bottom of the plot.

In summary, the mimic of an industrial scale-up process, validates the use of control circuits founded on biosensors for biosynthetic pathway intermediates coupled to expression of essential genes, to prolong and improve the productive lifespan of yeast cell factories for anabolic end-product formation.

## Discussion

Engineering microorganisms to produce valuable chemicals and fuels has the potential to enable the transition towards a greener and more sustainable bio-based economy (Nielsen and Keasling, 2016). However, the metabolic burden caused by engineering heterologous pathways for anabolic product formation in cell factories is often associated with reduced fitness and growth rate (Strucko et al., 2015; Wu et al., 2016). This ultimately imposes a selective pressure on engineered cells to reroute nutrients away from burdening non-essential pathways, or to delete these pathways completely, towards maximising cell proliferation (Rugbjerg et al., 2018a, 2018b).

In this study, the stability of a fitness-burdened yeast cell factory was characterised and compared with strains expressing two different control circuit designs founded on small-molecule biosensors coupled to essential gene expression. Based on the result of the sequential shake flasks experiments we determined that strains expressing control circuits outperformed the parental strain without a control circuit, as strains expressing control circuits enabled strains to remain productive for >55 generations in batch cultivations, compared to merely 25 generations observed for the parental strain (Figure 2, Figure 5). Moreover, comparing the performance of these strains in a fed-batch cultivation mimicking industrial setups, strains expressing control circuits accumulated up to 5-fold higher amounts of total pathway metabolites, including 72% and 92% higher end-product formation for VAC and PAC control circuits, respectively. This demonstrates that biosensors derived from prokaryotic small-molecule binding transcriptional regulators, can be successfully employed in yeast to expand the productive lifespan and control evolutionary drift of fitness-burdened cell factories, even when coupling essential gene expression with formation of product pathway intermediates.

Moreover, this study highlights how physiological, genetic, and bioinformatic approaches enable the engineering of control circuits founded on pathway intermediates. Specifically, by constructing single knockout gene deletions for all heterologous pathway steps, we were able to identify the key fitness-burdening pressure point in strains expressing the VG pathway. Next, by mining publicly available transcriptome and phenome datasets (Cherry et al., 2012; Regenberg et al., 2006), this study demonstrates successful selection criteria for identification of candidate essential genes, encoding a broad array of metabolic functions, which can be coupled to biosensors and thus enable selective growth based on small-molecule concentrations. With the increasing number of biosensors being developed, and demonstrations of fermentation-based manufacturing of valuable chemicals based on anabolic metabolism (Koch et al., 2019; Nielsen and Keasling, 2016), this combined approach should be applicable for stabilizing and optimizing many more bioprocesses in the future.

Having said this, this study not only exemplifies the error-modes of a fitness-burdening heterologous biosynthetic pathway, but also elucidates those observed in genetically-encoded control circuits, as those adopted in this study. First, by employing both transcriptional activators and repressors we observed that the PcaQ activator allowed for a wider range of essential genes to be used (Figure 4). Here, when replacing the native promoters with the PcaO-containing minimal *CYC1* promoter controlled by PcaQ, most of the tested essential genes were only able to sustain growth in strains producing PAC. On the contrary, when VanR was employed, all tested genes, with the exception of *GLN1*, showed similar growth upon VAC supplementation compared to control conditions. This is likely due to the lower OFF state of essential gene expression provided by the PAC control circuit founded on the minimal PcaO-containing *CYC1* promoter, highlighting the importance of biosensor output tuning when coupled to growth-based selection. Moreover, since many repressor-type biosensors established in *S. cerevisiae* rely on reporters with strong promoters to provide a readable output (Ambri et al., 2020; Dabirian et al., 2019; David et al., 2016; Hector and Mertens, 2017), this issue is believed to be a common concern for future strain stability efforts employing transcriptional repressors. Indeed, even though both types of control circuits lead to an increase in transcription in the presence of the ligand, inactivating mutations in the control circuits will lead to different outcomes depending on the aTF mode-of-action (D’Ambrosio and Jensen, 2017). With a repressor-based control circuit, any mutation limiting DNA-binding affinity will lock the expression of the actuating gene (e.g. an essential gene) in a quasi-ON state, just as deletion of non-native aTF repressor binding sites in output promoter is likely to do (Ambri et al., 2020). This quasi-ON state indeed proved to be sufficient for *GLN1* in this study (Figure 5B and Figure 6C). Contrastingly, if an activator-type aTF is mutated, the system will be locked in an OFF state, maintaining non-producing cells in a low-fitness stage. Thus, we recommend designs of control circuits based on short sequence-diverse synthetic promoters (Kotopka and Smolke, 2019; Redden and Alper, 2015) and activator-type aTFs to limit the rapid spreading of simple error-modes towards quasi-ON expression of output genes or homologous recombination events as observed for the “sniper-attack” on 30 bp VanO in the *TEF1*-based output promoter (Figure 6C).

Still, with meticulous characterization of the error-modes of cell factories and the input-output relationship connecting aTF biosensors with essential metabolic functions, control circuits are transforming biosensors from mere high-throughput screening technologies into an integral part of stabilizing bioprocesses, as already exemplified for bacterial cell factories (Rugbjerg et al., 2018b; Wang and Dunlop, 2019; Xiao et al., 2016). Ultimately, this will also enable understanding of the underlying mechanisms controlling population heterogeneity, and significantly contribute to development of new robust and cost-effective bio-based production processes.

## Methods

### Cultivation media and conditions

Chemically competent *Escherichia coli* DH5α strain was used as a host for cloning and plasmid propagation. The cells were cultivated at 37°C in 2xYT supplemented with 100 μg/mL ampicillin. The *Saccharomyces cerevisiae* strains used in this study were grown at 30°C and 250 rpm in three types of media: yeast extract peptone (YP) medium (10 g/L Bacto yeast extract and 10 g/L Bacto peptone), synthetic complete (SC) medium (6.7 g/L yeast nitrogen base without amino acids with appropriate drop-out medium supplement) and synthetic medium (SM) prepared as previously described (Mans et al., 2018). All three media were supplemented with glucose 20 g/L as carbon source unless otherwise specified. For media preparation, salts and water were sterilised in the autoclave at 120°C, before the complete mixed media were sterile filtered with 0.2 um filters.

### Plasmids and strains construction

All plasmids used in this study were assembled by USER™ (uracil-specific excision reagent) cloning (New England Biolabs). Biobricks constituting promoters, genes and other genetic elements required to assemble the plasmids were amplified by PCR using PhusionU polymerase (Thermo Fisher Scientific). *S. cerevisiae* strains were constructed by the lithium acetate/single-stranded carrier DNA/PEG method previously described (Gietz and Schiestl, 2007). The complete list of strains and plasmids used in this study is provided in Supplementary Tables S1 and S2.

### HPLC detection of extracellular metabolites

All metabolites of the vanillin β-glucoside pathway were analysed by HPLC using the Dionex Ultimate 3000 HPLC (Thermo Fisher Scientific), coupled with the Supelco Discovery HS F5-3 HPLC column (150 ×2.1 mm × 3 µm) (Sigma Aldrich). Mobile phase A consisted of 10 mM ammonium formate, pH 3 while mobile phase B consisted of acetonitrile. The elution profile was as follows: 5% of solvent B for 0.5 min and increased linearly to 60% B over 5 min. The gradient was increased to 90% B over 0.5 min and kept at this condition for 2 min. Finally, returned to 5% B and equilibrated until 10 min. The flow rate was set at 0.7mL/minute while the column was held at 30°C and the metabolites were detected using the UV diodide detector DAD-3000 Diode Array Detector set at 260, 277, 304 and 210 nm. The samples were prepared as previously described (Strucko et al., 2017). Shortly, 1 mL of yeast culture and 1 mL of 96% ethanol were mixed. The solution was then centrifuged at 12000g for 2 minutes. The supernatant was then collected and stored at −20°C until it was measured on the HPLC.

### Biosensor design and promoter replacement

The bidirectional VanR design used to replace essential genes promoters is composed of the *ADH1* terminator, VanR from *Caulobacter crescentus*, the *PGK1* promoter driving the expression of VanR and the engineered TEF1p containing two VanO sequences separated by the Eco47III restriction site. The design as described was used to replace the 150bp sequence upstream of the selected essential genes. The PcaQ biosensor design is composed of two parts: PcaQ, under the control of the strong TDH3 promoter and the reporter module composed of the truncated CYC1 promoter containing the PcaQ binding site, both previously described (Ambri et al., 2020). The complete list of the gRNAs used in this study can be found in Supplementary Table S3. Each gRNA sequence was then combined with a backbone carrying the pRNR2-Cas9-CYC1t cassette to assemble an all-in-one CRISPR plasmid, by USER™ (uracil-specific excision reagent) cloning (New England Biolabs), as previously described (D’Ambrosio et al., 2020). In order to maximise the transformation efficiency the plasmid were used to transform yeast strains where one additional copy of TEF1p-Cas9-CYC1t was already present in the genome within the EasyClone site X-4 site (Jensen et al., 2014).

### Growth profiler analysis

Cell growth was evaluated using the Growth Profiler 960 (Enzyscreen B.V., The Netherlands) at 30°C and 200 rpm. Prior to measurement, *S. cerevisiae* strains, with the exception of strains carrying the PcaQ biosensor design, were inoculated in 0.5 mL of synthetic medium in 96-format polypropylene deep-well plates, and grown overnight at 30°C and 300 rpm. The cells were then diluted 1:100 in fresh synthetic medium. Because of the longer lag phase or inability to grow in SM medium (Supplementary Figure S6), the strains where the native essential gene promoter was replaced with the PcaQ biosensor design were initially inoculated in rich YPD medium and grown overnight. Next, the strains were diluted 1:10 in synthetic medium and cultured overnight. Finally, the strains were diluted 1:100 in fresh synthetic medium to a final volume of 150 μL and the cell suspension was transferred to a 96-half deepwell plate (Enzyscreen B.V., The Netherlands). In order to convert the G-values provided by the instrument into OD600 values we generated a calibration curve by measuring the G-value of samples with known OD.

### Flow cytometry measurements

Single cell fluorescence was evaluated using a Becton Dickinson LSR FORTESSA with a blue 488 nm laser. Prior to measurement, *S. cerevisiae* strains were grown overnight in mineral medium at 30°C and 300 rpm in 96-format polypropylene deep-well plates. Next, the cells were diluted 1:50 in fresh mineral medium and were cultivated for 20 hours. Finally, cells were diluted 1:20 in PBS to arrest cell growth and the fluorescence was measured by flow cytometry. For each sample 10,000 single-cell events were recorded.

### Whole genome sequencing

The samples selected for whole genome sequence were inoculated from glycerol stock in rich YPD medium and cultured overnight. The genomic DNA was extracted by using the Quick-DNA Fungal/Bacterial Miniprep Kit (Zymoresearch) following the provided protocol. DNA libraries were then prepared using a Kapa Hyper Prep Library Prep Kit (Roche) and sequenced by Illumina MiSeq. Data mapping was performed against the CEN.PK113-7D genome (Salazar et al., 2017) where additional expression cassettes relative to vanillin-β-glucoside biosynthetic pathway genes and control circuits were previously added. Data processing and chromosome copy number variation determinations were performed as previously described (Nijkamp et al., 2012; Verhoeven et al., 2017).

### Fed-batch cultivations

The bioreactor cultures were performed in 1 L bioreactors (Biostat Q Plus, Sartorius, Gottingen, Germany) and started with a batch phase using 400 mL synthetic medium containing 20 g/L of glucose and 5 mL Antifoam 204 (Sigma A6426). The temperature was maintained at 30°C and the pH was kept constant at 5.0 by dropwise addition of a 10M KOH solution. The cultures were sparged with pressurized air at a flow rate of 500 mL/min and dissolved oxygen levels were maintained above 50% by controlling the stirrer speed. The off-gas CO_2_ and O_2_ was monitored throughout the cultivation (Prima BT Mass Spectrometer, Thermo Fisher Scientific) and the data was acquired by the Lucullus software (Securecell AG, Switzerland). After carbon depletion, indicated by a rapid rise in dissolved oxygen levels, the fed-batch phase was automatically initiated (Supplementary Figure S9). During the fed-batch phase, a 2x concentrated SM solution containing 200 g/L glucose and Antifoam (1mL/L), was used. The initial feed rate was set to 2 mL h^-1^, resulting in a growth rate of 0.05 h^-1^. Throughout the fed-batch phase, the feed rate was continuously increased with a rate of 0.05 h^-1^ to maintain a constant growth rate. Samples were taken manually at defined time points during the batch and fed-batch phase and metabolite consumption and formation were determined via HPLC analysis.

## Acknowledgements

This work was supported by the Novo Nordisk Foundation and by the European Union’s Horizon 2020 research and innovation programme under the Marie Sklodowska-Curie action PAcMEN (grant agreement No. 722287). Finally, we would like to thank Dr. Tomas Strucko for sharing with us the parental *S. cerevisiae* VG strain used in this study.

## Author contributions

RM, PR, MOAS, JDK and MKJ conceived the study. VD, ED, RDB, FA and RM conducted all experimental work related to strain designs and construction, MvdB performed all next-generation sequencing analysis, while JtH and SS conducted all fermentation. VD, RM, and MKJ wrote the manuscript.

