## Supplementary information for "Regulatory control circuits for stabilizing long-term anabolic product formation in yeast"

**Supplementary Table S1: List of *S. cerevisiae* strains used in this study.**

| Strain ID | Background strain | Genotype | Reference/ Resource |
|---|---|---|---|
| VG | | MATα MAL2-8C SUC2 XII2(pTEF1-HsOMT, pPGK1-UGT) XII1(pTEF1-PPT, pPGK1-ACAR) XII5(pTEF1-3DSD) Δbgl1::loxP Δadh6::KanMX | (Hansen et al., 2009) |
| yRV_01 | VG | MATα MAL2-8C SUC2 XII2(pTEF1-HsOMT, pPGK1-UGT) XII1(pTEF1-PPT, pPGK1-ACAR) XII5(pTEF1-3DSD), XI3(pPGK1-VanR, pTEF1(2xVanO)-yeGFP) Δbgl1::loxP Δadh6::KanMX | This study |
| yRV_02 | VG | MATα MAL2-8C SUC2 XII2(pTEF1-HsOMT, pPGK1-UGT) XII1(pTEF1-PPT, pPGK1-ACAR) Δbgl1::loxP Δadh6::KanMX Δ3DSD::hphMX | This study |
| yRV_03 | VG | MATα MAL2-8C SUC2 XII2(pTEF1-HsOMT, pPGK1-UGT) XII1(pTEF1-PPT) XII5(pTEF1-3DSD) Δbgl1::loxP Δadh6::KanMX ΔACAR::hphMX | This study |
| yRV_04 | VG | MATα MAL2-8C SUC2 XII2(pTEF1-HsOMT, pPGK1-UGT) XII1( pPGK1-ACAR) XII5(pTEF1-3DSD) Δbgl1::loxP Δadh6::KanMX ΔPPT::hphMX | This study |
| yRV_05 | VG | MATα MAL2-8C SUC2 XII2(pTEF1-HsOMT) XII1(pTEF1-PPT, pPGK1-ACAR) XII5(pTEF1-3DSD) Δbgl1::loxP Δadh6::KanMX ΔUGT::hphMX | This study |
| yRV_06 | VG | MATα MAL2-8C SUC2 XII2(pPGK1-UGT) XII1(pTEF1-PPT, pPGK1-ACAR) XII5(pTEF1-3DSD) Δbgl1::loxP Δadh6::KanMX ΔhsOMT::hphMX | This study |
| CEN. PK113-7D | | Mat a MAL2-8c SUC2 URA3 HIS3 LEU2 TRP1 | (Nijkamp et al., 2012) |
| yRV_07 | CEN.PK113-7D | Mat a MAL2-8c SUC2 URA3 HIS3 LEU2 TRP1 X4(TEF1p-Cas9) | This study |
| yRV_08 | yRV_07 | Mat a MAL2-8c SUC2 URA3 HIS3 LEU2 TRP1 X4(TEF1p-Cas9) pFOL2(pPGK1-VanR, pTEF1(2xVanO))-FOL2 | This study |

| | | | |
|---|---|---|---|
| yRV_09 | yRV_07 | Mat a MAL2-8c SUC2 URA3 HIS3 LEU2 TRP1 X4(TEF1p-Cas9) pDUT1(pPGK1-VanR, pTEF1(2xVanO))-DUT1 | This study |
| yRV_10 | yRV_07 | Mat a MAL2-8c SUC2 URA3 HIS3 LEU2 TRP1 X4(TEF1p-Cas9) pCYS3(pPGK1-VanR, pTEF1(2xVanO))-CYS3 | This study |
| yRV_11 | yRV_07 | Mat a MAL2-8c SUC2 URA3 HIS3 LEU2 TRP1 X4(TEF1p-Cas9) pCHO1(pPGK1-VanR, pTEF1(2xVanO))-CHO1 | This study |
| yRV_12 | yRV_07 | Mat a MAL2-8c SUC2 URA3 HIS3 LEU2 TRP1 X4(TEF1p-Cas9) pGUK1(pPGK1-VanR, pTEF1(2xVanO))-GUK1 | This study |
| yRV_13 | yRV_07 | Mat a MAL2-8c SUC2 URA3 HIS3 LEU2 TRP1 X4(TEF1p-Cas9) pGLN1(pPGK1-VanR, pTEF1(2xVanO))-GLN1 | This study |
| yRV_14 | yRV_07 | Mat a MAL2-8c SUC2 URA3 HIS3 LEU2 TRP1 X4(TEF1p-Cas9) pFAS1(pPGK1-VanR, pTEF1(2xVanO))-FAS1 | This study |
| yRV_15 | yRV_07 | Mat a MAL2-8c SUC2 URA3 HIS3 LEU2 TRP1 X4(TEF1p-Cas9) pURA2(pPGK1-VanR, pTEF1(2xVanO))-URA2 | This study |
| yRV_16 | yRV_07 | Mat a MAL2-8c SUC2 URA3 HIS3 LEU2 TRP1 X4(TEF1p-Cas9) pOLE1(pPGK1-VanR, pTEF1(2xVanO))-OLE1 | This study |
| yRV_17 | yRV_07 | Mat a MAL2-8c SUC2 URA3 HIS3 LEU2 TRP1 X4(TEF1p-Cas9) pARO2(pPGK1-VanR, pTEF1(2xVanO))-ARO2 | This study |
| yRV_18 | VG | MATα MAL2-8C SUC2 XII2(pTEF1-HsOMT, pPGK1-UGT) XII1(pTEF1-PPT, pPGK1-ACAR) XII5(pTEF1-3DSD) X4(TEF1p-Cas9) Δbgl1::loxP Δadh6::KanMX | This study |
| yRV_19 | yRV_02 | MATα MAL2-8C SUC2 XII2(pTEF1-HsOMT, pPGK1-UGT) XII1(pTEF1-PPT, pPGK1-ACAR) X4(TEF1p-Cas9) Δbgl1::loxP Δadh6::KanMX Δ3DSD::hphMX | This study |

| | | | |
|---|---|---|---|
| yRV_20 | yRV_18 | MATα MAL2-8C SUC2 XII2(pTEF1-HsOMT, pPGK1-UGT) XII1(pTEF1-PPT, pPGK1-ACAR) Δbgl1::loxP Δadh6::KanMX X4(TEF1p-Cas9) pFOL2(209bp_CYC1(PcaO))-FOL2 | This study |
| yRV_21 | yRV_18 | MATα MAL2-8C SUC2 XII2(pTEF1-HsOMT, pPGK1-UGT) XII1(pTEF1-PPT, pPGK1-ACAR) Δbgl1::loxP Δadh6::KanMX X4(TEF1p-Cas9) pDUT1(209bp_CYC1(PcaO))-DUT1 | This study |
| yRV_22 | yRV_18 | MATα MAL2-8C SUC2 XII2(pTEF1-HsOMT, pPGK1-UGT) XII1(pTEF1-PPT, pPGK1-ACAR) Δbgl1::loxP Δadh6::KanMX X4(TEF1p-Cas9) pCYS3(209bp_CYC1(PcaO))-CYS3 | This study |
| yRV_23 | yRV_18 | MATα MAL2-8C SUC2 XII2(pTEF1-HsOMT, pPGK1-UGT) XII1(pTEF1-PPT, pPGK1-ACAR) Δbgl1::loxP Δadh6::KanMX X4(TEF1p-Cas9) pCHO1(209bp_CYC1(PcaO))-CHO1 | This study |
| yRV_24 | yRV_18 | MATα MAL2-8C SUC2 XII2(pTEF1-HsOMT, pPGK1-UGT) XII1(pTEF1-PPT, pPGK1-ACAR) Δbgl1::loxP Δadh6::KanMX X4(TEF1p-Cas9) pGUK1(209bp_CYC1(PcaO))-GUK1 | This study |
| yRV_25 | yRV_18 | MATα MAL2-8C SUC2 XII2(pTEF1-HsOMT, pPGK1-UGT) XII1(pTEF1-PPT, pPGK1-ACAR) Δbgl1::loxP Δadh6::KanMX X4(TEF1p-Cas9) pGLN1(209bp_CYC1(PcaO))-GLN1 | This study |
| yRV_26 | yRV_18 | MATα MAL2-8C SUC2 XII2(pTEF1-HsOMT, pPGK1-UGT) XII1(pTEF1-PPT, pPGK1-ACAR) Δbgl1::loxP Δadh6::KanMX X4(TEF1p-Cas9) pFAS1(209bp_CYC1(PcaO))-FAS1 | This study |
| yRV_27 | yRV_18 | MATα MAL2-8C SUC2 XII2(pTEF1-HsOMT, pPGK1-UGT) XII1(pTEF1-PPT, pPGK1-ACAR) Δbgl1::loxP Δadh6::KanMX X4(TEF1p-Cas9) pURA2(209bp_CYC1(PcaO))-URA2 | This study |
| yRV_28 | yRV_18 | MATα MAL2-8C SUC2 XII2(pTEF1-HsOMT, pPGK1-UGT) XII1(pTEF1-PPT, pPGK1-ACAR) Δbgl1::loxP | This study |

| | | | |
|---|---|---|---|
| | | Δadh6::KanMX X4(TEF1p-Cas9) pOLE1(209bp_CYC1(PcaO))-OLE1 | |
| yRV_29 | yRV_18 | MATα MAL2-8C SUC2 XII2(pTEF1-HsOMT, pPGK1-UGT) XII1(pTEF1-PPT, pPGK1-ACAR) Δbgl1::loxP Δadh6::KanMX X4(TEF1p-Cas9) pARO2(209bp_CYC1(PcaO))-ARO2 | This study |
| yRV_30 | yRV_19 | MATα MAL2-8C SUC2 XII2(pTEF1-HsOMT, pPGK1-UGT) XII1(pTEF1-PPT, pPGK1-ACAR) Δbgl1::loxP Δadh6::KanMX Δ3DSD::hphMX X4(TEF1p-Cas9) pFOL2(209bp_CYC1(PcaO))-FOL2 | This study |
| yRV_31 | yRV_19 | MATα MAL2-8C SUC2 XII2(pTEF1-HsOMT, pPGK1-UGT) XII1(pTEF1-PPT, pPGK1-ACAR) Δbgl1::loxP Δadh6::KanMX Δ3DSD::hphMX X4(TEF1p-Cas9) pDUT1(209bp_CYC1(PcaO))-DUT1 | This study |
| yRV_32 | yRV_19 | MATα MAL2-8C SUC2 XII2(pTEF1-HsOMT, pPGK1-UGT) XII1(pTEF1-PPT, pPGK1-ACAR) Δbgl1::loxP Δadh6::KanMX Δ3DSD::hphMX X4(TEF1p-Cas9) pCYS3(209bp_CYC1(PcaO))-CYS3 | This study |
| yRV_33 | yRV_19 | MATα MAL2-8C SUC2 XII2(pTEF1-HsOMT, pPGK1-UGT) XII1(pTEF1-PPT, pPGK1-ACAR) Δbgl1::loxP Δadh6::KanMX Δ3DSD::hphMX X4(TEF1p-Cas9) pCHO1(209bp_CYC1(PcaO))-CHO1 | This study |
| yRV_34 | yRV_19 | MATα MAL2-8C SUC2 XII2(pTEF1-HsOMT, pPGK1-UGT) XII1(pTEF1-PPT, pPGK1-ACAR) Δbgl1::loxP Δadh6::KanMX Δ3DSD::hphMX X4(TEF1p-Cas9) pGUK1(209bp_CYC1(PcaO))-GUK1 | This study |
| yRV_35 | yRV_19 | MATα MAL2-8C SUC2 XII2(pTEF1-HsOMT, pPGK1-UGT) XII1(pTEF1-PPT, pPGK1-ACAR) Δbgl1::loxP Δadh6::KanMX Δ3DSD::hphMX X4(TEF1p-Cas9) pGLN1(209bp_CYC1(PcaO))-GLN1 | This study |
| yRV_36 | yRV_19 | MATα MAL2-8C SUC2 XII2(pTEF1-HsOMT, pPGK1-UGT) XII1(pTEF1-PPT, pPGK1-ACAR) Δbgl1::loxP Δadh6::KanMX Δ3DSD::hphMX X4(TEF1p-Cas9) pFAS1(209bp_CYC1(PcaO))-FAS1 | This study |

| | | | |
|---|---|---|---|
| yRV_37 | yRV_19 | MATα MAL2-8C SUC2 XII2(pTEF1-HsOMT, pPGK1-UGT) XII1(pTEF1-PPT, pPGK1-ACAR) Δbgl1::loxP Δadh6::KanMX Δ3DSD::hphMX X4(TEF1p-Cas9) pURA2(209bp_CYC1(PcaO))-URA2 | This study |
| yRV_38 | yRV_19 | MATα MAL2-8C SUC2 XII2(pTEF1-HsOMT, pPGK1-UGT) XII1(pTEF1-PPT, pPGK1-ACAR) Δbgl1::loxP Δadh6::KanMX Δ3DSD::hphMX X4(TEF1p-Cas9) pOLE1(209bp_CYC1(PcaO))-OLE1 | This study |
| yRV_39 | yRV_19 | MATα MAL2-8C SUC2 XII2(pTEF1-HsOMT, pPGK1-UGT) XII1(pTEF1-PPT, pPGK1-ACAR) Δbgl1::loxP Δadh6::KanMX Δ3DSD::hphMX X4(TEF1p-Cas9) pARO2(209bp_CYC1(PcaO))-ARO2 | This study |
| yRV_40 | yRV_18 | MATα MAL2-8C SUC2 XII2(pTEF1-HsOMT, pPGK1-UGT) XII1(pTEF1-PPT, pPGK1-ACAR) Δbgl1::loxP Δadh6::KanMX X4(TEF1p-Cas9) pGLN1(pPGK1-VanR, pTEF1(2xVanO))-GLN1 | This study |
| yRV_41 | yRV_02 | MATα MAL2-8C SUC2 XII2(pTEF1-HsOMT, pPGK1-UGT) XII1(pTEF1-PPT, pPGK1-ACAR) XI3(pPGK1-VanR, pTEF1(2xVanO)-yeGFP) Δbgl1::loxP Δadh6::KanMX Δ3DSD::hphMX | This study |

**Supplementary Table S2: List of plasmids used in this study.**

| Plasmid ID | Parental plasmid | Description | Reference/ Source |
|---|---|---|---|
| pCFB390 | | pXI-3-loxP-KlURA3 | (Jensen et al., 2014) |
| pGC_06 | pCFB390 | pXI-3-loxP-KlURA3-VanR<-PGK1-pTEF1(2xVanO) ->yeGFP | This study |

| | | | |
|---|---|---|---|
| pTS-33 | | pX-3-LoxP-KlLEU2-TDH3p->PcaQ | (Skjoedt et al., 2016) |
| pFA197 | | pXII-4-LoxP-SpHIS5-209bp-pCYC1(PcaO-108)->yeGFP | (Ambri et al., 2020) |
| pCFB1767 | | pRS414-TEF1p-Cas9-CYC1t | (DiCarlo et al., 2013) |
| pTajak177 | | NatMx, pRNR2-Cas9-CYC1t | (D'Ambrosio et al., 2020) |
| pRV_01 | pTajak177 | NatMx, pRNR2-Cas9-CYC1t, FOL2 gRNA cassette | This study |
| pRV_02 | pTajak177 | NatMx, pRNR2-Cas9-CYC1t, DUT1 gRNA cassette | This study |
| pRV_03 | pTajak177 | NatMx, pRNR2-Cas9-CYC1t, CYS3 gRNA cassette | This study |
| pRV_04 | pTajak177 | NatMx, pRNR2-Cas9-CYC1t, CHO1 gRNA cassette | This study |
| pRV_05 | pTajak177 | NatMx, pRNR2-Cas9-CYC1t, GUK1 gRNA cassette | This study |
| pRV_06 | pTajak177 | NatMx, pRNR2-Cas9-CYC1t, GLN1 gRNA cassette | This study |
| pRV_07 | pTajak177 | NatMx, pRNR2-Cas9-CYC1t, FAS1 gRNA cassette | This study |
| pRV_08 | pTajak177 | NatMx, pRNR2-Cas9-CYC1t, URA2 gRNA cassette | This study |
| pRV_09 | pTajak177 | NatMx, pRNR2-Cas9-CYC1t, OLE1 gRNA cassette | This study |
| pRV_10 | pTajak177 | NatMx, pRNR2-Cas9-CYC1t, ARO2 gRNA cassette | This study |

**Supplementary Table S3: gRNA cassettes used in this study.** The gRNA sequence is highlighted (yellow) while primer binding regions are lowercase

| **Essential gene promoter** | **Sequence** |
|---|---|
| *FOL2* | agggaacaaaagctggagctTCTTTGAAAAGATAATGTATGATTATGCT<br>TTCACTCATATTTATACAGAAACTTGATGTTTTCTTTCGAGTA<br>TATACAAGGTGATTACATGTACGTTTGAAGTACAACTCTAGA<br>TTTTGTAGTGCCCTCTTGGGCTAGCGGTAAAGGTGCGCATTTT<br>TTCACACCCTACAATGTTCTGTTCAAAAGATTTTGGTCAAACG<br>CTGTAGAAGTGAAAGTTGGTGCGCATGTTTCGGCGTTCGAAA<br>CTTCTCCGCAGTGAAAGATAAATGATCCGCACGCACAATCAC<br>GATTGGTTTTAGAGCTAGAAATAGCAAGTTAAAATAAGGCTA<br>GTCCGTTATCAACTTGAAAAAGTGGCACCGAGTCGGTGGTGC<br>TTTTTTTGTTTTTTATGTCTtcgagtcatgtaattagtta |
| *DUT1* | agggaacaaaagctggagctTCTTTGAAAAGATAATGTATGATTATGCT<br>TTCACTCATATTTATACAGAAACTTGATGTTTTCTTTCGAGTA<br>TATACAAGGTGATTACATGTACGTTTGAAGTACAACTCTAGA<br>TTTTGTAGTGCCCTCTTGGGCTAGCGGTAAAGGTGCGCATTTT<br>TTCACACCCTACAATGTTCTGTTCAAAAGATTTTGGTCAAACG<br>CTGTAGAAGTGAAAGTTGGTGCGCATGTTTCGGCGTTCGAAA<br>CTTCTCCGCAGTGAAAGATAAATGATCAGTTGTTTCTACTTAT<br>TAAAGTTTTAGAGCTAGAAATAGCAAGTTAAAATAAGGCTAG<br>TCCGTTATCAACTTGAAAAAGTGGCACCGAGTCGGTGGTGCT<br>TTTTTTGTTTTTTATGTCTtcgagtcatgtaattagtta |
| *CYS3* | agggaacaaaagctggagctTCTTTGAAAAGATAATGTATGATTATGCT<br>TTCACTCATATTTATACAGAAACTTGATGTTTTCTTTCGAGTA<br>TATACAAGGTGATTACATGTACGTTTGAAGTACAACTCTAGA<br>TTTTGTAGTGCCCTCTTGGGCTAGCGGTAAAGGTGCGCATTTT<br>TTCACACCCTACAATGTTCTGTTCAAAAGATTTTGGTCAAACG<br>CTGTAGAAGTGAAAGTTGGTGCGCATGTTTCGGCGTTCGAAA<br>CTTCTCCGCAGTGAAAGATAAATGATCGAATTTTGAAAGTAC<br>AATTGGTTTTAGAGCTAGAAATAGCAAGTTAAAATAAGGCTA<br>GTCCGTTATCAACTTGAAAAAGTGGCACCGAGTCGGTGGTGC<br>TTTTTTTGTTTTTTATGTCTtcgagtcatgtaattagtta |
| *CHO1* | agggaacaaaagctggagctTCTTTGAAAAGATAATGTATGATTATGCT<br>TTCACTCATATTTATACAGAAACTTGATGTTTTCTTTCGAGTA |

| | |
|---|---|
| | TATACAAGGTGATTACATGTACGTTTGAAGTACAACTCTAGA TTTTGTAGTGCCCTCTTGGGCTAGCGGTAAAGGTGCGCATTTT TTCACACCCTACAATGTTCTGTTCAAAAGATTTTGGTCAAACG CTGTAGAAGTGAAAGTTGGTGCGCATGTTTCGGCGTTCGAAA CTTCTCCGCAGTGAAAGATAAATGATCATAATGTTTCCTATA AAATAGTTTTAGAGCTAGAAATAGCAAGTTAAAATAAGGCTA GTCCGTTATCAACTTGAAAAAGTGGCACCGAGTCGGTGGTGC TTTTTTTGTTTTTTATGTCTtcgagtcatgtaattagtta |
| *GUK1* | agggaacaaaagctggagctTCTTTGAAAAGATAATGTATGATTATGCT TTCACTCATATTTATACAGAAACTTGATGTTTTCTTTCGAGTA TATACAAGGTGATTACATGTACGTTTGAAGTACAACTCTAGA TTTTGTAGTGCCCTCTTGGGCTAGCGGTAAAGGTGCGCATTTT TTCACACCCTACAATGTTCTGTTCAAAAGATTTTGGTCAAACG CTGTAGAAGTGAAAGTTGGTGCGCATGTTTCGGCGTTCGAAA CTTCTCCGCAGTGAAAGATAAATGATCTTGATAACTACAGTT TACTTGTTTTAGAGCTAGAAATAGCAAGTTAAAATAAGGCTA GTCCGTTATCAACTTGAAAAAGTGGCACCGAGTCGGTGGTGC TTTTTTTGTTTTTTATGTCTtcgagtcatgtaattagtta |
| *GLN1* | agggaacaaaagctggagctTCTTTGAAAAGATAATGTATGATTATGCT TTCACTCATATTTATACAGAAACTTGATGTTTTCTTTCGAGTA TATACAAGGTGATTACATGTACGTTTGAAGTACAACTCTAGA TTTTGTAGTGCCCTCTTGGGCTAGCGGTAAAGGTGCGCATTTT TTCACACCCTACAATGTTCTGTTCAAAAGATTTTGGTCAAACG CTGTAGAAGTGAAAGTTGGTGCGCATGTTTCGGCGTTCGAAA CTTCTCCGCAGTGAAAGATAAATGATCGTTCTGTCTTTGTTTT CGTTGTTTTAGAGCTAGAAATAGCAAGTTAAAATAAGGCTAG TCCGTTATCAACTTGAAAAAGTGGCACCGAGTCGGTGGTGCT TTTTTTGTTTTTTATGTCTtcgagtcatgtaattagtta |
| *FAS1* | agggaacaaaagctggagctTCTTTGAAAAGATAATGTATGATTATGCT TTCACTCATATTTATACAGAAACTTGATGTTTTCTTTCGAGTA TATACAAGGTGATTACATGTACGTTTGAAGTACAACTCTAGA TTTTGTAGTGCCCTCTTGGGCTAGCGGTAAAGGTGCGCATTTT TTCACACCCTACAATGTTCTGTTCAAAAGATTTTGGTCAAACG CTGTAGAAGTGAAAGTTGGTGCGCATGTTTCGGCGTTCGAAA CTTCTCCGCAGTGAAAGATAAATGATCGGAAAATCAGATTAT AGACTGTTTTAGAGCTAGAAATAGCAAGTTAAAATAAGGCTA |

| | |
|---|---|
| | GTCCGTTATCAACTTGAAAAAGTGGCACCGAGTCGGTGGTGC TTTTTTTGTTTTTTATGTCTtcgagtcatgtaattagtta |
| *URA2* | agggaacaaaagctggagctTCTTTGAAAAGATAATGTATGATTATGCT TTCACTCATATTTATACAGAAACTTGATGTTTTCTTTCGAGTA TATACAAGGTGATTACATGTACGTTTGAAGTACAACTCTAGA TTTTGTAGTGCCCTCTTGGGCTAGCGGTAAAGGTGCGCATTTT TTCACACCCTACAATGTTCTGTTCAAAAGATTTTGGTCAAACG CTGTAGAAGTGAAAGTTGGTGCGCATGTTTCGGCGTTCGAAA CTTCTCCGCAGTGAAAGATAAATGATCTTGTTATATTCTATTA GGTAGTTTTAGAGCTAGAAATAGCAAGTTAAAATAAGGCTAG TCCGTTATCAACTTGAAAAAGTGGCACCGAGTCGGTGGTGCT TTTTTTGTTTTTTATGTCTtcgagtcatgtaattagtta |
| *OLE1* | agggaacaaaagctggagctTCTTTGAAAAGATAATGTATGATTATGCT TTCACTCATATTTATACAGAAACTTGATGTTTTCTTTCGAGTA TATACAAGGTGATTACATGTACGTTTGAAGTACAACTCTAGA TTTTGTAGTGCCCTCTTGGGCTAGCGGTAAAGGTGCGCATTTT TTCACACCCTACAATGTTCTGTTCAAAAGATTTTGGTCAAACG CTGTAGAAGTGAAAGTTGGTGCGCATGTTTCGGCGTTCGAAA CTTCTCCGCAGTGAAAGATAAATGATCATCATAGTAATAGAT AGTTGGTTTTAGAGCTAGAAATAGCAAGTTAAAATAAGGCTA GTCCGTTATCAACTTGAAAAAGTGGCACCGAGTCGGTGGTGC TTTTTTTGTTTTTTATGTCTtcgagtcatgtaattagtta |
| *ARO2* | agggaacaaaagctggagctTCTTTGAAAAGATAATGTATGATTATGCT TTCACTCATATTTATACAGAAACTTGATGTTTTCTTTCGAGTA TATACAAGGTGATTACATGTACGTTTGAAGTACAACTCTAGA TTTTGTAGTGCCCTCTTGGGCTAGCGGTAAAGGTGCGCATTTT TTCACACCCTACAATGTTCTGTTCAAAAGATTTTGGTCAAACG CTGTAGAAGTGAAAGTTGGTGCGCATGTTTCGGCGTTCGAAA CTTCTCCGCAGTGAAAGATAAATGATCAAAATAGTATCATAG CACAGGTTTTAGAGCTAGAAATAGCAAGTTAAAATAAGGCTA GTCCGTTATCAACTTGAAAAAGTGGCACCGAGTCGGTGGTGC TTTTTTTGTTTTTTATGTCTtcgagtcatgtaattagtta |

**Supplementary Table S4. Lists of essential genes according to the six imposed selection criteria**. Gene lists are colour-coded in purple (amino acid metabolism), light yellow (nucleotide metabolism), grey (fatty acid metabolism) and light blue (cofactor and vitamin metabolism).

*See separate Suppl. table S4 file*

**Supplementary Table S5. Amino acid abundance in VG pathway enzymes and yeast biomass.** The abundance of each amino acid in the pathway (PW) is expressed as a percentage of the occurring amino acid over the total number of amino acids in the pathway related proteins. The BioM% represents the average amino acid composition of the yeast biomass while the difference between the two values is represented in the P-B column.

| **Amino acid** | | **PW %** | **BioM %** | **P-B** |
|---|---|---|---|---|
| Q | Gln | 3,98% | 7,76% | -3,77% |
| K | Lys | 3,46% | 6,58% | -3,13% |
| G | Gly | 6,39% | 8,91% | -2,52% |
| N | Asn | 2,48% | 4,65% | -2,17% |
| E | Glu | 6,69% | 7,76% | -1,07% |
| I | Ile | 5,15% | 5,90% | -0,75% |
| T | Thr | 5,19% | 5,58% | -0,40% |
| F | Phe | 3,72% | 3,77% | -0,05% |
| V | Val | 7,63% | 7,35% | 0,28% |
| S | Ser | 5,71% | 5,34% | 0,37% |

| | | | | |
|---|---|---|---|---|
| H | His | 2,56% | 1,93% | 0,62% |
| W | Trp | 1,28% | 0,65% | 0,63% |
| A | Ala | 10,48% | 9,79% | 0,69% |
| M | Met | 1,84% | 1,14% | 0,70% |
| Y | Tyr | 3,04% | 1,96% | 1,08% |
| P | Pro | 5,37% | 4,23% | 1,14% |
| C | Cys | 1,47% | 0,14% | 1,33% |
| D | Asp | 6,50% | 4,65% | 1,85% |
| R | Arg | 6,35% | 3,87% | 2,48% |
| L | Leu | 10,71% | 8,03% | 2,68% |

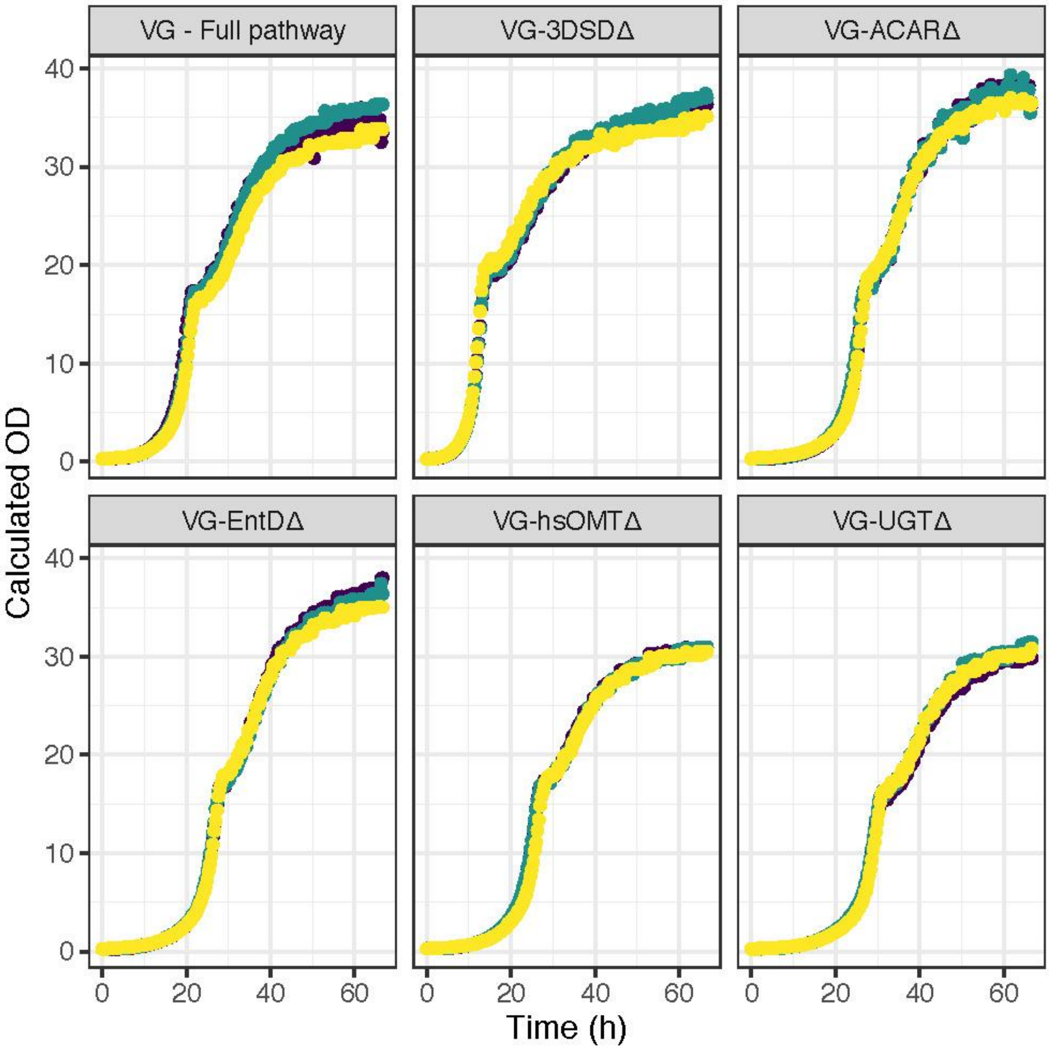


**Supplementary Figure S1.** Growth profiles (n=3) of single knockouts strains and VG strain carrying the complete vanillin-β-glucoside biosynthetic pathway.

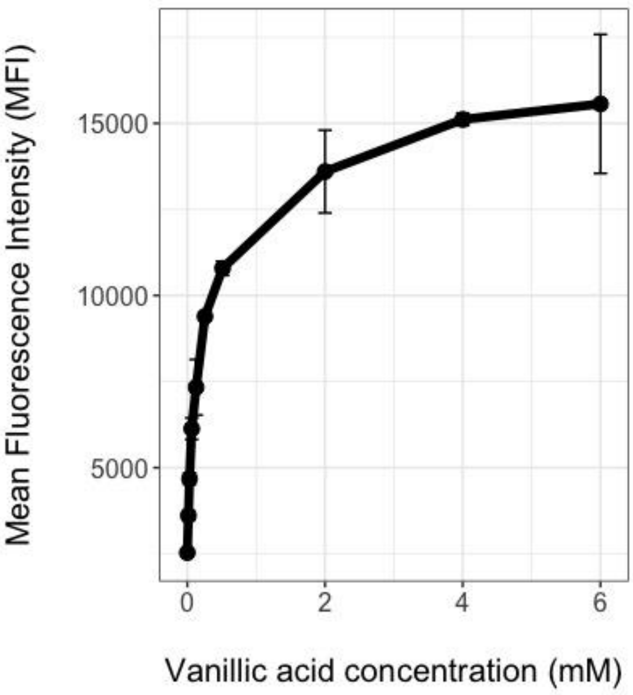


**Supplementary Figure S2. VanR biosensor design characterisation**. Characterisation of the response of the VanR biosensor design in response to increasing concentrations of vanillic acid. The points represent the average MFI of three (n=3) replicates. Error bars represent mean ± standard deviation from three (n=3) biological replicates.

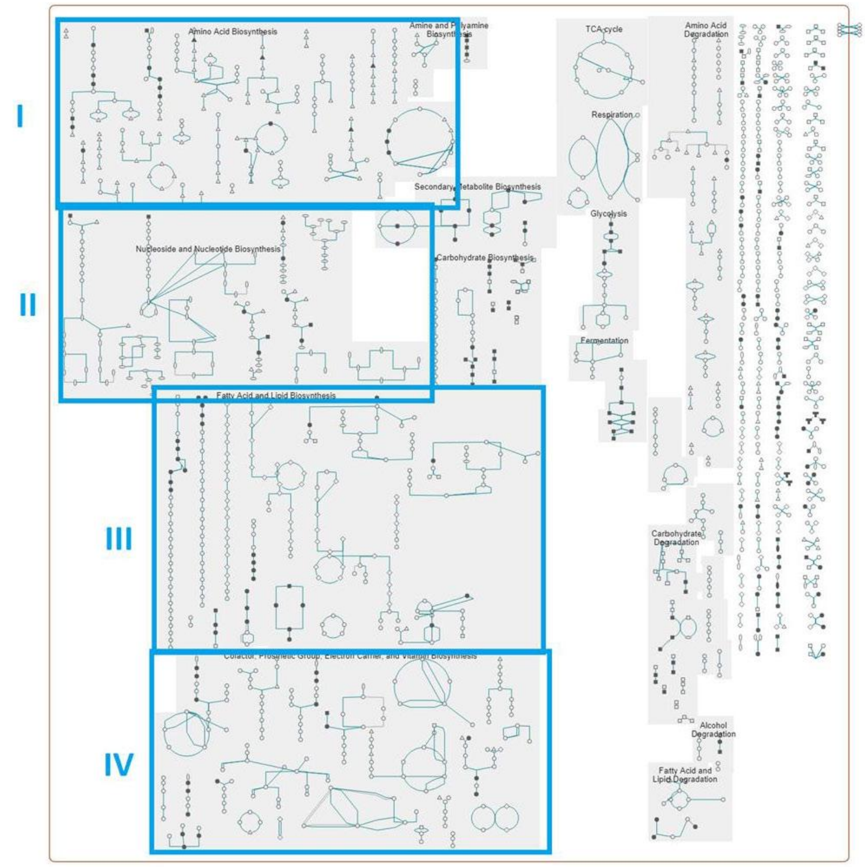


**Supplementary Figure S3. Metabolic network of *S. cerevisiae.*** Metabolic network of *Saccharomyces cerevisiae*, with the 4 main classes of biosynthetic reactions included in the selected essential genes highlighted in blue boxes. The classes of reactions include biosynthesis of amino acids, biosynthesis of nucleosides and nucleotides (II), biosynthesis of fatty acids and lipids (III), and biosynthesis of cofactors and vitamins (IV)

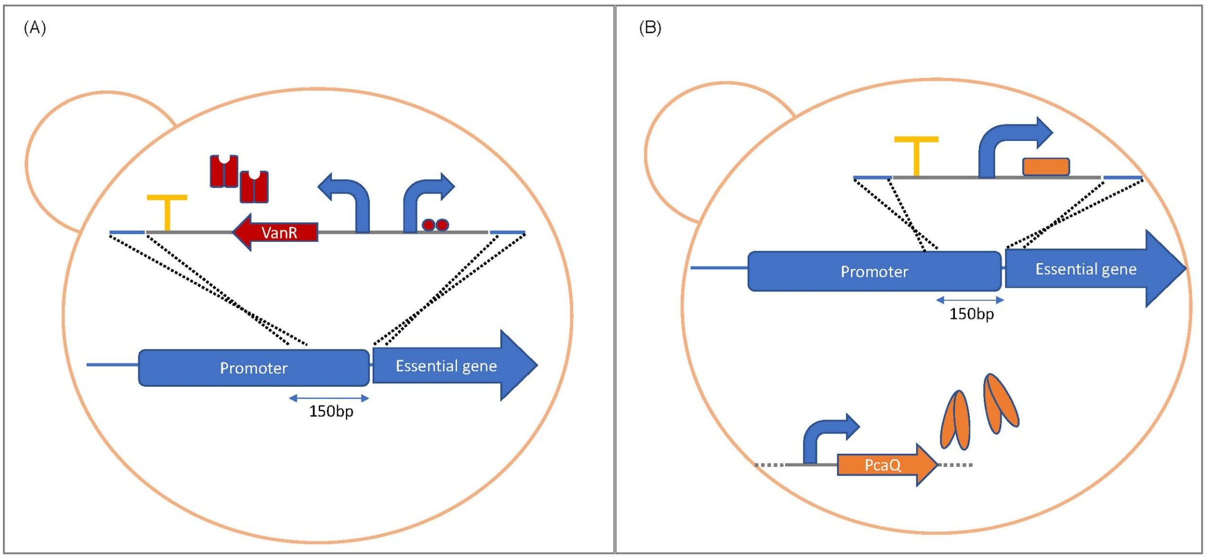


**Supplementary Figure S4. Schematic representation of the essential gene promoter replacement.** (**A**) Schematic representation of the promoter replacement by the bi-directional VanR biosensor design. The integration linear fragment is composed as follows: CYC1t (yellow T), *VANR* gene (red arrow) controlled by the PGK1p (blue arrow) and synthetic TEF1p (blue arrow) containing two copies of the VanO sequence (red circles) separated by the Eco47III restriction site. VanR dimers are represented in red. (**B**) Schematic representation of the promoter replacement by the PcaQ biosensor design. The integration linear fragment is composed as follows: CYC1t (yellow T), synthetic truncated CYC1p (209bp_CYC1p) containing one copy of the PcaO site (orange box). PcaQ dimers are depicted in orange. The PcaQ cassette was previously integrated in EasyClone site X-3. (A-B) Both the biosensor designs are designed to replace the 150bp sequence upstream of the essential gene sequence.

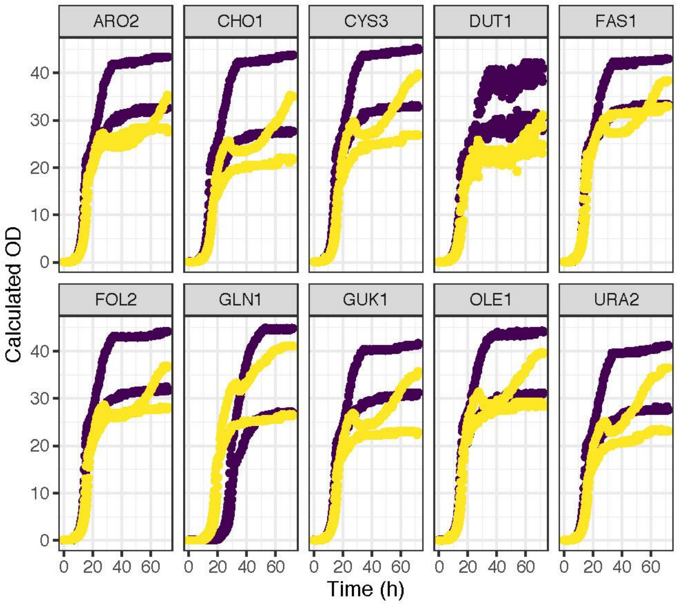


**Supplementary Figure S5. VAC control circuit characterisation.** Growth profiles (n=2) of wild type CEN.PK strains carrying the VAC control circuit design in the presence (yellow) or absence (purple) of 2mM vanillic acid.

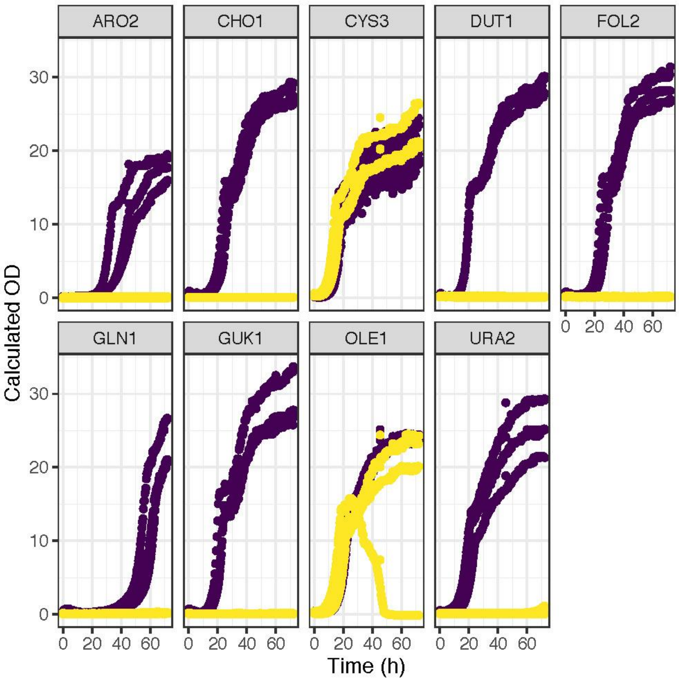


**Supplementary Figure S6. PAC control circuit characterisation.** Growth profiles (n=3) of VG pathway strains (purple) and VG-3DSDΔ strains (yellow) carrying the PAC control circuit design. It was not possible to identify a colony having the *FAS1* promoter replaced by the PAC control circuit.

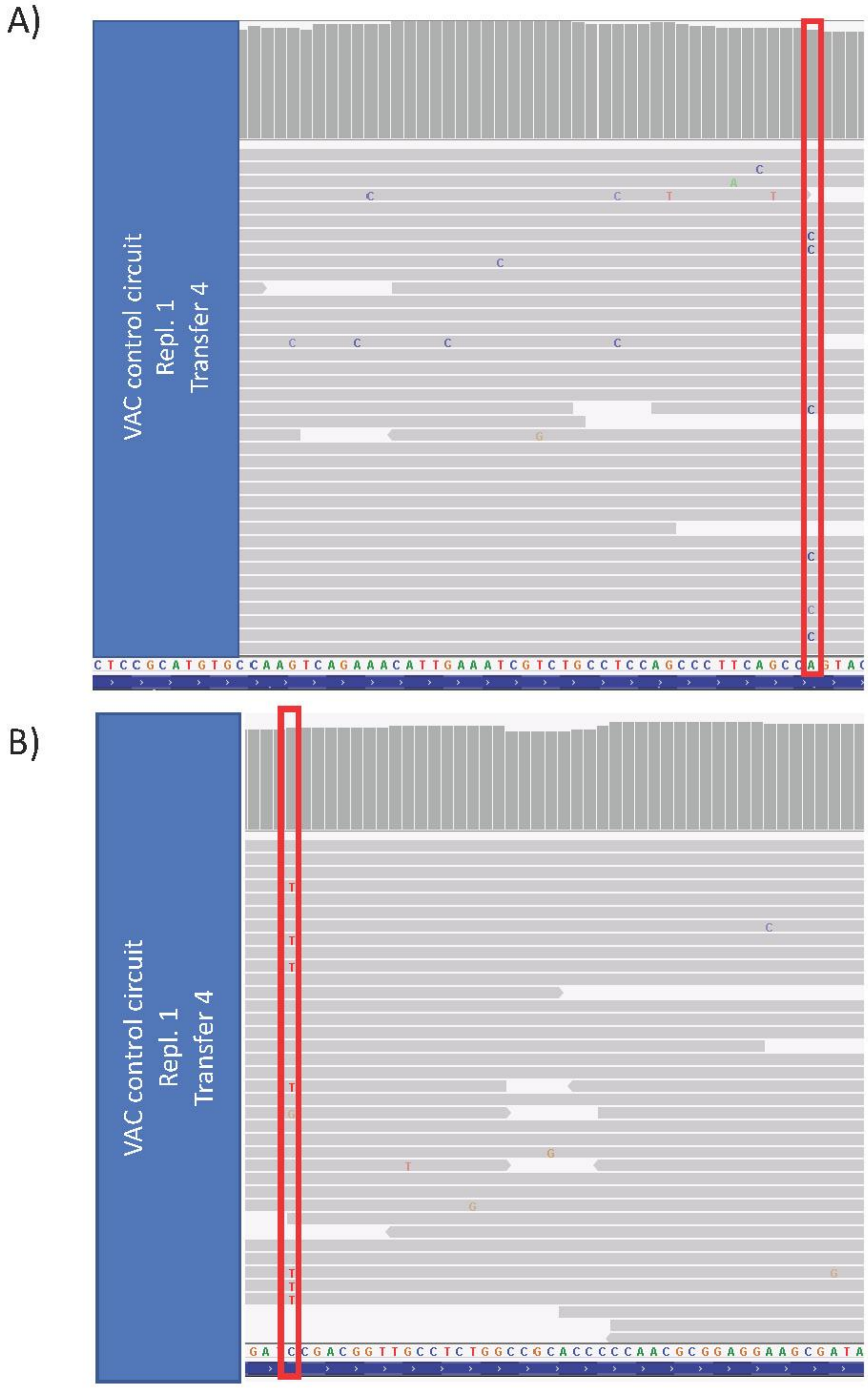


**Supplementary Figure S7. Manually identified mutations in *3DSD* gene for VAC control circuit replicate 1.** (**A-B**) Genome viewer zooms on the *3DSD* gene for the VAC control circuit strain replicate 1 following transfer 4. (**A**) Identification of the non-synonymous mutation Q91P by manual inspection of the *3DSD* gene reads. Blue segment represents *3DSD* ORF while the red segment highlights the mutated bases. (**B**) Identification of the non-synonymous mutation P208S by manual inspection of the *3DSD* gene reads. Blue segment represents *3DSD* ORF while the red segment highlights the mutated bases.

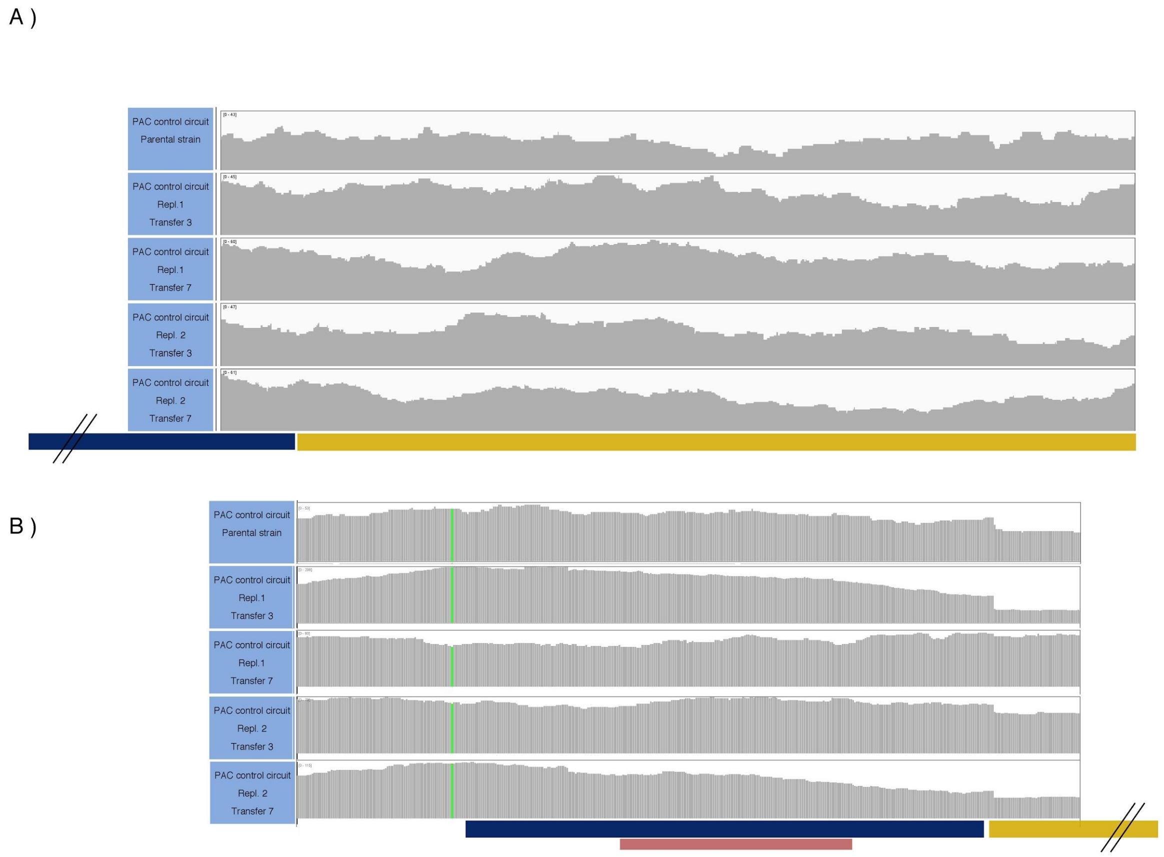


**Supplementary Figure S8. PAC control circuit population sequencing.**

(**A**) Genome-viewer zooms on the *3DSD* gene of the two replicate PAC control circuit strains at transfer 3 and 7 and of the parental strain. Above the bar plots for read coverage, the genomic layout of the *3DSD* gene expression unit is indicated with the *TEF1* presented in blue and the *3DSD* gene in yellow. (**B**) Genome-viewer zoom on the synthetic PcaO-containing truncated *CYC1* promoter of the PAC control circuit strain. The PcaO operator site is marked with a red box. Above the bar plot for read coverage, the genomic layout of the *GLN1* gene expression unit is indicated with the *209bpCYC1-VanO* promoter presented in blue and the *GLN1* gene in yellow. For (B) green bar represents a SNP compared to the CEN.PK113-7D reference genome.

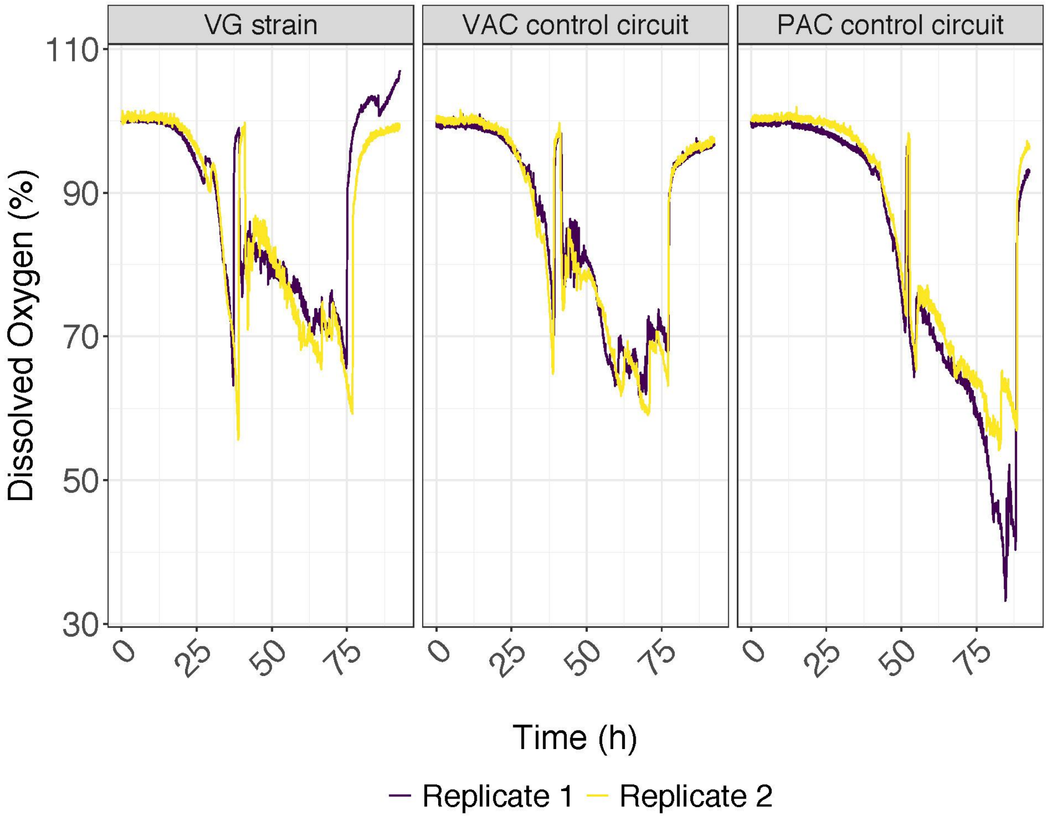


**Supplementary Figure S9. Dissolved oxygen levels during batch and fed-batch cultivations.** (**A-C**) Dissolved oxygen levels for the parental VG strain (left panel), and strains expressing VAC (middle panel) and PAC (right panel) control circuits. A spike in the oxygen levels indicates carbon depletion, upon which the fed-batch phase was started. Each line represents the dissolved oxygen levels for one biological replicate.

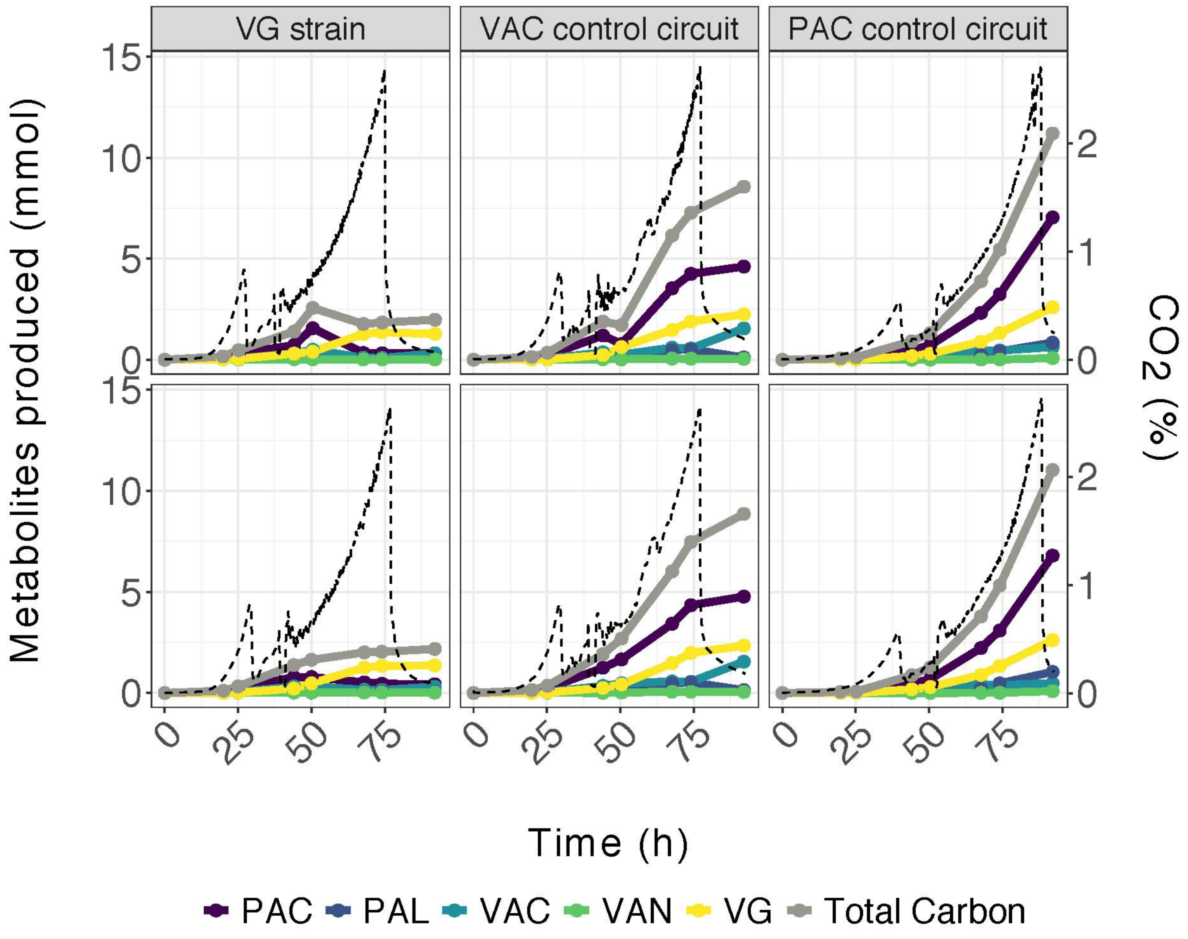


**Supplementary Figure S10. Extracellular metabolites characterisation during fed-batch cultivations.**

Extracellular metabolite characterization of the parental VG strain (left panel), and strains expressing VAC (middle panel) and PAC (right panel) control circuits. Metabolite production is expressed in mmol of VG pathway metabolites produced during the batch and fed-batch phases. The values were corrected for the medium volume used for HPLC sampling. Dashed black lines represent off-gas $CO_2$ (percent) while coloured lines indicate accumulation of individual VG pathway metabolites during the cultivation (Time, h) according to colour coding indicated at the bottom of the plot.
